# Aging reprograms the response to chronic stress

**DOI:** 10.64898/2026.02.16.702827

**Authors:** Gregor Stein, Mihail Ivilinov Todorov, Lisa Lange, Konstantin Riege, Emilio Cirri, Iqra Hussain, Martina Korfei, Nadine Pömpner, Christian A. Hübner, Karl Lenhard Rudolph, Farida Hellal, Ali Ertürk, Steve Hoffmann, Olivia Engmann

**Affiliations:** Institute for Biochemistry and Biophysics, Friedrich-Schiller-University Jena, Germany; Institute for Intelligent Biotechnologies (iBIO), Helmholtz Center Munich, Neuherberg, Germany; Institute for Stroke and Dementia Research (ISD), University Hospital, LMU Munich, Munich, Germany; Computational Biology Group, Leibniz Institute on Aging - Fritz Lipmann Institute (FLI), Beutenbergstraße 11, 07745, Jena, Germany; Proteomics Core Facility, Leibniz Institute on Aging – Fritz Lipmann Institute (FLI), Beutenbergstraße 11, 07745 Jena, Germany; Institute of Human Genetics, Jena University Hospital, Am Klinikum 1, F2E20, 07747, Jena, Germany; Leibniz Institute on Aging - Fritz Lipmann Institute (FLI), Beutenbergstraße 11, 07745 Jena, Germany; Munich Cluster for Systems Neurology (SyNergy), Munich, Germany; Deep Piction, Munich, Germany; School of Medicine, Koç University, İstanbul, Turkey

**Keywords:** Aging, chronic stress, depression, resilience, animal model, angiogenesis, proteomics, Omics

## Abstract

Chronic stress is thought to accelerate brain aging. We find this to be true in the brains of young mice, but reversed in the old. Using chronic variable stress in young (2-month) and aged (24-month) mice, we show that aged animals perceive stress physiologically but exhibit stress-responses that differ from those of young mice on behavioral, synaptic, and molecular levels. Multi-Omics profiling of prefrontal cortex and nucleus accumbens and 3D vasculature measurements reveals that stress in aged mice activates angiogenic programs that oppose aging-related patterns. Our results demonstrate that stress cannot be universally conceptualized as an aging accelerator, but instead engages age-specific programs with opposite directionality in young and old animals.

## Introduction

Chronic stress is a main risk factor for depression, the leading cause of disability worldwide ^1,2^. Accordingly, stress-based animal models are widely used to study molecular mechanisms underlying depression and to identify therapeutic targets. However, these models typically rely on early-life or young-adult animals ^3–5^. Therefore, the question how aging reshapes the neurobiological response to chronic stress, is currently unresolved.

The highest prevalence of depression is found between 18-25 years of age ^6,7^. Depression and affective disorders are the main cause for hospitalization at age 15-24 years in Germany before pregnancy and intoxication ^8^. In contrast, depression in the elderly is more commonly associated with comorbidities such as chronic medical conditions, cognitive impairment, and bereavement, rather than chronic stress ^9,10^. Older individuals may also benefit from protective psychological factors such as socio-emotional selectivity, raising the possibility that resilience to chronic stress increases with age ^10^. Together, these epidemiological data imply that aged individuals may engage stress-response programs distinct from those in young cohorts.

In parallel, recent work has proposed the “accelerated aging” hypothesis, which states that chronic stress and depression drive age-related brain changes. Supporting evidence includes accelerated age-dependent volume loss in striatal reward regions, including the nucleus accumbens (NAc), in patients with depression ^11^. Moreover, the research conducted by Zannas and colleagues highlights the role of transcriptional regulation by the glucocorticoid receptor, telomere shortening, and epigenetic mechanisms, including DNA methylation, in mediating the effects of stress on aging ^12^. However, this framework is derived largely from young populations, leaving it unknown whether stress involves similar aging-associated programs in the aged brain. Here, we assess how pathways of aging and stress interact by using the chronic variable stress (CVS) mouse model, which allows for sex-specific investigations. We integrate behavioral and physiological phenotyping with transcriptomics, proteomics, dendritic spine morphology, and three-dimensional vascular imaging, including two stress-relevant brain regions NAc and prefrontal cortex (PFC) as well as the liver.

In line with the accelerated aging hypothesis, we find that chronic stress overlaps with aging-associated molecular signatures in young mice. In contrast, in aged mice, stress is accompanied with reduced depression-like phenotypes and age-associated signatures, including molecular reversal, synaptic and vascular remodeling. These data revise the accelerated aging framework and establish that aging re-programs the direction of stress-induced changes in the brain.

## Methods

Further information can be found in the *extended methods*.

### Animals and Licenses

Mice were housed following the ethical guidelines of the Thüringer Landesamt für Verbraucherschutz (TLV). Experiments were conducted under animal licenses UKJ-18-037, FSU-22-003 and FSU-25-002 (Germany), which comply with the EU Directive 2010/63/EU guidelines for animal experiments. Experiments with genetically modified organisms were performed according to S1 regulations according to the GenTAufzV. C57Bl/6J mice were housed and bred in the animal facility (FZL) of the Universitätsklinikum Jena, Germany, the Fritz-Lipmann Institute for Aging Research and in the BIZ animal facility of Friedrich-Schiller-University Jena, Germany, or purchased from Janvier Labs (Saint Berthevin Cedex, France). Both sexes were used as stated. Young mice were 8-12 weeks old at the beginning of stress induction, while aged mice were 98-113 weeks, or approximately 2 years, old. They were housed in a 12L:12D light cycle. Aged mice included ex-breeders, reflecting child rearing as a physiological component of aging. In order to avoid bias towards the fittest individuals, aged mice were also included in the study if they showed minor health issues such as barbering, above or below average weight. These individuals were monitored closely and balanced across groups.

### AAVs and stereotaxic surgery

Bilateral stereotaxic surgery into the PFC was essentially performed as described ^13^. A GFP-expressing virus (pAAV.1-CAG-GFP (#37825, Addgene) was utilized at a concentrations of app. 6 x 10^11^ (GFP) particles * ml^-1^.

### Behavioral tests and CVS

Behavioral testing and chronic variable stress (CVS) were performed as described ^5,14^. Mice underwent 21 days of CVS consisting of tube restraint, tail suspension, or mild foot shocks administered in a semi-random order, followed by assessment in the forced swim and splash tests during the active (dark) phase. Animal health and body weight were scored throughout the procedure.

### Dendritic spine analysis

The analysis was based on detecting the AAVs’ GFP-fluorophores. Photos of 40 µm paraformaldehyde-fixed brain sections were taken with a Zeiss LSM 880 confocal microscope using the AiryScan method. Maximum intensity projections were obtained using Zen Black and Zen Blue software and analyzed in NeuronStudio (CNIC, Mount Sinai School of Medicine). The total density of spines, proportions of neck-containing and stubby spines as well as the cumulative neck length were determined in Graph Prism.

### Next-generation RNA-sequencing

RNA was purified by resuspension in Trizol and chloroform-precipitation. RNA was washed in isopropanol and 75% Ethanol as described ^15^.Transcriptomics was performed as previously described ^16^. Poly(A)-enriched mRNAs from 500 ng total RNA per sample were processed using a stranded library preparation protocol and subsequently amplified for sequencing. Reads were quality-assessed, preprocessed and subsquently aligned against the mouse reference genome mm10, followed by UMI-based removal of PCR duplicates, gene expression quantification and differential testing. Data tables are provided via the Gene Expression Omnibus (GEO) database together with raw sequencing data under accession number GSE317903.

### Proteomics

Tissue proteins were extracted under denaturing conditions, reduced and alkylated, and processed by S-trap–based tryptic digestion before peptide cleanup and loading onto Evotips for LC–MS analysis. Peptides were separated by Evosep One chromatography and analyzed by DIA on an Orbitrap Exploris 480, followed by directDIA processing in Spectronaut using a mouse SwissProt database for identification and label-free quantification. Differential protein abundance was assessed by precursor-level statistics with Benjamini–Hochberg correction. Mass spectrometry proteomics data have been deposited to the ProteomeXchange Consortium via the MassIVE partner repository, and they are accessible with the identifiers MassIVE (MSV000100662_reviewer (PFC), MSV000100634_reviewer (NAc) and MSV000100635_reviewer (liver)).

### Tissue labeling, optical clearing, and light-sheet imaging of brain vasculature

Animals were anesthetized with ketamine/xylazine (100/16 mg/kg, i.p.), and upon complete loss of nociceptive reflexes, 50 µl Lycopersicon esculentum (Tomato) lectin–DyLight 594 was injected into the left ventricle to label the perfused vasculature for 60 s. Brains were then extracted and post-fixed in 4% PFA for 12 h, followed by optical clearing using an adapted 3DISCO protocol involving graded tetrahydrofuran dehydration, dichloromethane delipidation, and refractive index matching in benzyl alcohol/benzyl benzoate (1:2). Cleared brains were imaged on a LaVision UltraII light-sheet microscope using 2x and 4x objectives, with DyLight 594 fluorescence detected at 580/25 nm excitation and 625/30 nm emission. To minimize defocus, a five-step sequential focal shift was applied per plane, with bidirectional illumination and z-steps of 32.5 µm (2x) or 3 µm (4x). Vascular networks were segmented and quantified from 4x datasets using the VesSAP pipeline, extracting vessel length density, bifurcation density, and vessel radius, with all measurements corrected for clearing-induced tissue shrinkage.

### Statistics

Analyses were performed using GraphPad Prism with tests selected based on data distribution and experimental design, including non-parametric tests as well as parametric like One- and Two-way ANOVA with appropriate post-hoc corrections. Transcriptomic, proteomic, morphological, and vascular analyses were conducted in independent cohorts, with behavioral and imaging analyses performed blinded to condition and processed by independent experimenters or core facilities to minimize bias. Overlap between molecular datasets was assessed using rank–rank hypergeometric overlap (RRHO) analysis with Benjamini-Hochberg correction.

## Results

### Aging reverses behavioral and synaptic stress responses

To determine age-effects on the stress response, young and aged mice underwent 21 days of CVS (vs. naïve controls). Behavioral testing was performed 24 h after the final stress exposure.

In young mice, CVS induced depression-like behaviors, including altered immobility in the forced swim test and grooming in the splash test (**Fig. 1A-C**; **Fig. S1**) as previously observed ^13,16^. In contrast, CVS significantly reduced immobility time in aged mice compared to naïve mice of the same age, suggesting a reversal of stress-induced escape behavior with age (**Fig. 1B**). Grooming in the splash test was not affected by CVS in aged mice (**Fig. 1C**). Importantly, CVS induced comparable body-weight loss in young and aged mice, showing that aged mice can sense and physiologically react to stress (**Fig. 1D**, **Fig. S1**).

**Fig. 1:**
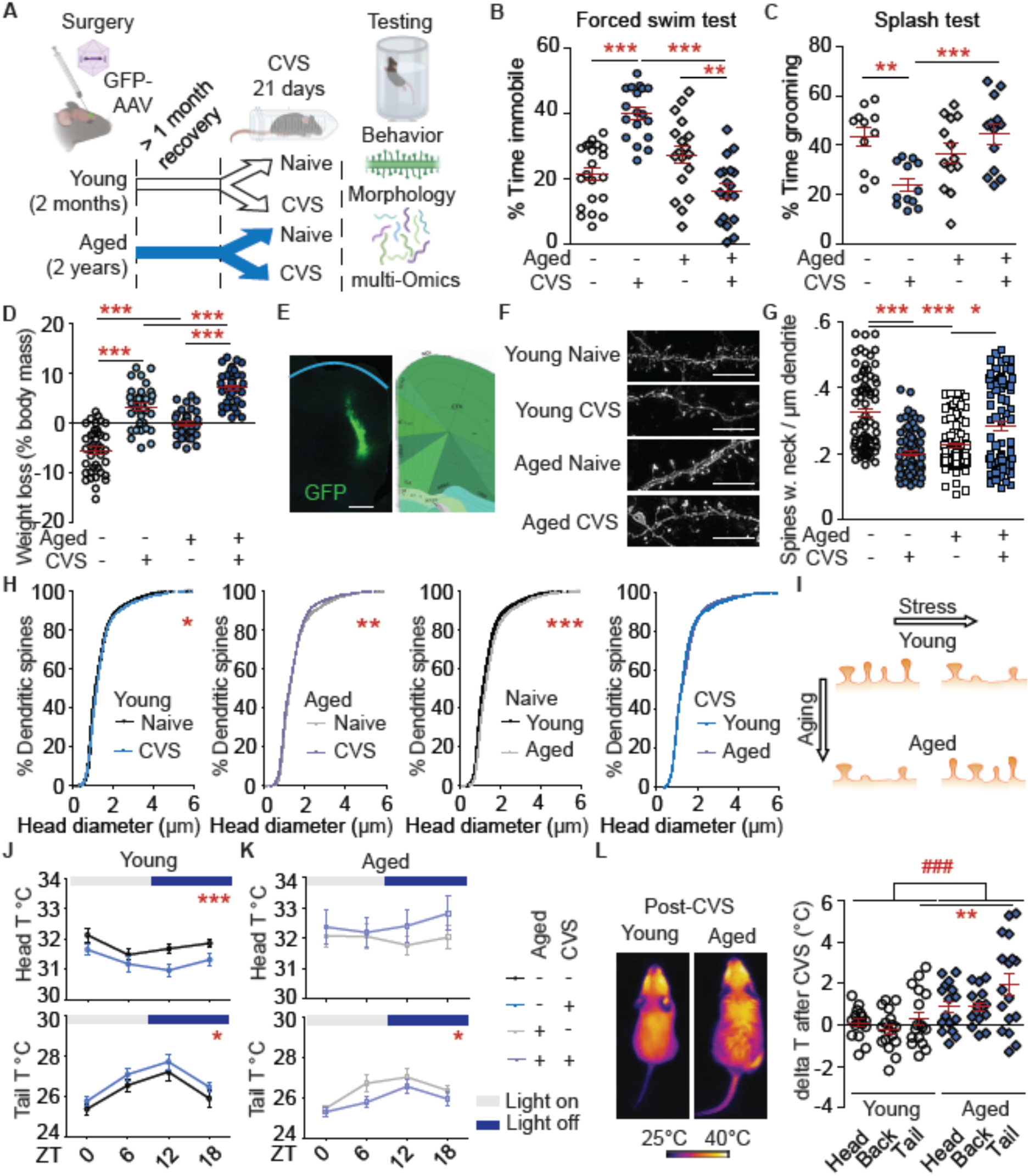
Aging affects the stress response on behavioral and morphological levels. **A**) Experimental design. **B**-**D**, **G**) Two-way ANOVA & Bonferroni *post hoc* test. Independent data points are plotted and means **±** s.e.m. are shown. **B**) Stress-induced immobility time in the forced swim test is increased in young mice but reduced in aged mice. **C**) Grooming in the splash test is reduced by CVS in young but not aged mice. **D**) The body weight is reduced by stress in both, young and aged mice. **E**) Overview of viral injection into the PFC (right part was generated using the Allen Brain atlas). Scale bar: 400 µm. **F**) Representative dendrites. Scale bar 10 µm. **G**) CVS reduces neck-containing spines in young mice. **H**) The cumulative head diameter is increased by CVS in young mice and by aging, while it is reduced by CVS in aged mice. For better visibility, the x-axis is arbitrarily set to 0-6 µm. Spines with a larger diameter were however included in the analysis. **I**) Summary of spine changes after stress and aging. **J**-**L**) The body temperature in young and aged mice were measured around the circadian clock (**J**, **K**) and directly before and after the last stress induction (**L**). **J**) In young mice, stress reduces the core body temperature and increases the peripheral temperature across the circadian cycle. **K**) In aged mice, stress increases the peripheral temperature across the circadian cycle. **L**) Stress induction on day 21 acutely increases the body temperature in aged mice more than in young ones. The reference time point just before stress induction refers to the ZT12 time point. Sketches were made with biorender.com. For this figure, both sexes were pooled. Sex-specific analyses and detailed statistics can be found in the supplemental material. ZT – Zeitgeber time.

We next assessed dendritic spine morphology in the PFC, a correlate of chronic stress and antidepressant action in mice ^13,16^ and humans ^17^. As expected ^13,18^, CVS in young mice and aging in naïve mice both reduced the density of neck-containing (thin and mushroom) spines and increased the cumulative head diameter, suggesting a loss of spines with smaller head sizes. In line with the behavioral data, stress showed an inverse effect on spines in aged mice. The density of neck-containing spines was increased and the head diameter reduced, suggesting a gain of spines with smaller head diameters (**Fig. 1E-I**; **Fig. S1**).

Physiological readouts similarly identified an age-dependent divergence of stress responses. In young mice, CVS induced circadian changes in body temperature as previously described ^19,20^. These included reduced head and back temperature, and increased tail temperature, suggesting a possible change in peripheral vascular tone. These stress responses were absent in aged mice, which instead showed a reduction in tail temperature following CVS (**Fig. 1J, K**; **Fig. S2**). The overall body temperature did not differ across ages (**Fig. S2**). CVS evoked stronger acute temperature changes in aged mice directly after stress induction, further demonstrating that aged mice are able to sense and respond to stress (**Fig. 1L**; **Fig. S2**). Thus, the altered behavioral, synaptic and thermal responses observed in aged mice do not reflect impaired stress sensing but rather a qualitative reprogramming of stress-response pathways with age.

### Stress elicits distinct molecular programs in young and aged brains

To define the molecular architecture of age-dependent stress responses, we performed sex-specific transcriptomics and proteomics profiling on two main centers of the stress response, the PFC and NAc ^5,21^ (**Tables S1-S12**). Across sexes and brain regions, stress in both ages induced an overall downregulation of mRNA transcripts, whereas the net direction of protein regulation was more balanced, suggesting partial uncoupling of transcriptional and translational stress responses (**Table S1**). In the PFC, aging itself showed a slight trend of upregulation within significantly differentially expressed genes (DEGs) as well as proteins (DEPs) across sexes and brain regions, suggesting regulatory mechanisms other than stress responses (**Fig. 2A**, **Table S1**).

**Fig. 2:**
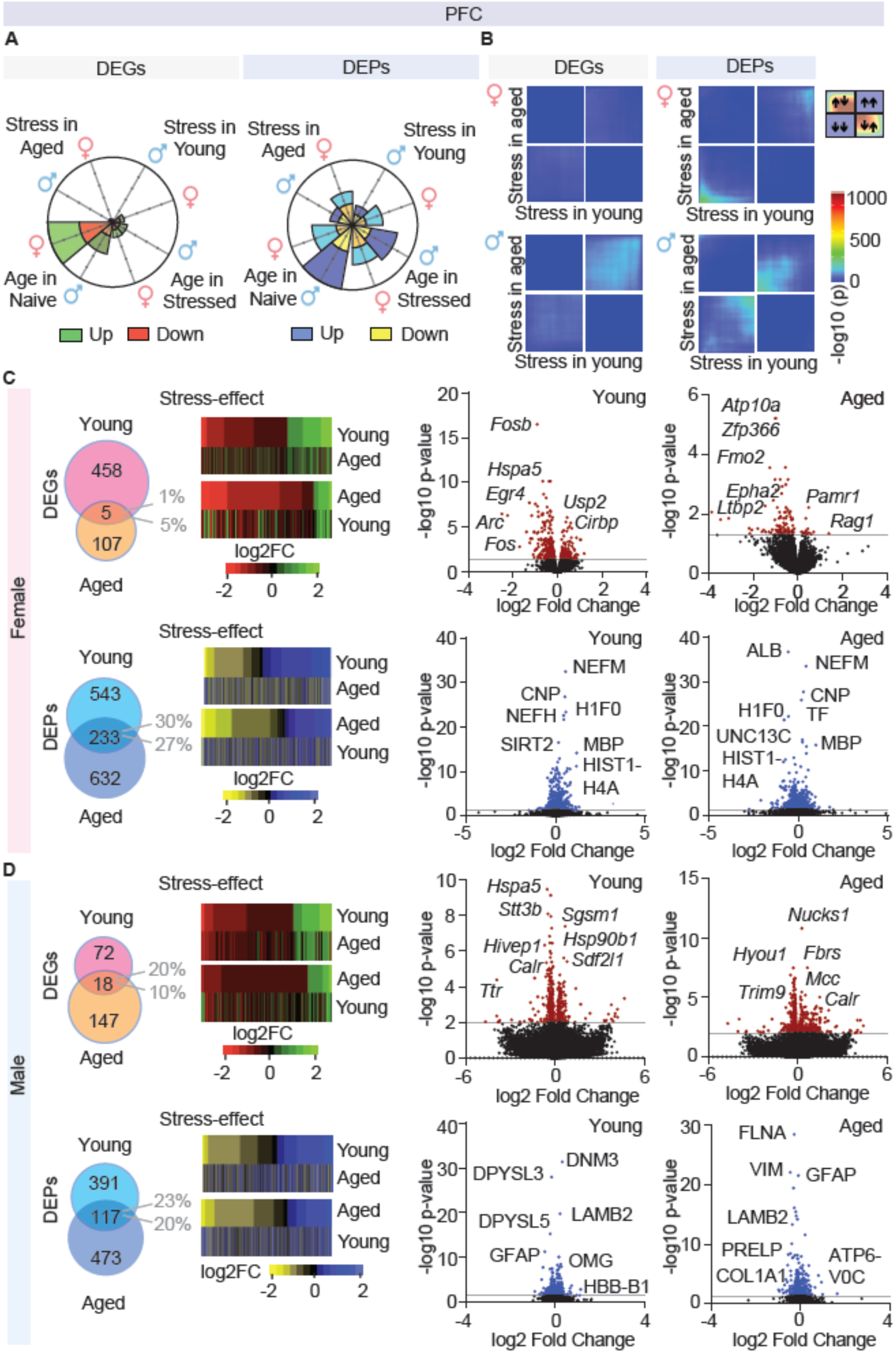
Little overlap of stress effects across age groups in the PFC. **A**) Petal plots show the total numbers of up- and downregulated DEGs and DEPs. **B**) RRHO plots show little association between stress effects in young versus aged mice across cohorts. **C**, **D**) Venn diagrams and heatmaps demonstrate extent and direction of overlapping DEGs and DEPs that are regulated by stress across age-groups. Heatmaps show significant stress-effects in one age group (first row), complemented by fold-changes of identical set within the second age group, including non-significant changes. Therefore, heatmaps were generated in which each group was on top as in ^5^. Heatmaps reveal a modest to no association of stress effects across age groups. Volcano plots depict top altered molecules by log10 p-value. Labels are selected for most strongly altered and representative gene products. **C**) Females. **D**) Males.

Rank-rank hypergeometric (RRHO) plots depict a weak concordance between stress-regulated DEGs and DEPs in young and aged mice across the PFC (**Fig. 2B**). Consistently, on average only a small percentage of stress-regulated DEGs and DEPs overlap between young and aged mice across sexes and brain regions with minimal directional concordance between stress-effects in young versus aged mice (**Fig. 2C, D**). The same patterns were observed in the NAc (**Fig. 3**) and in Pearson correlations (**Table S2**).

**Fig. 3:**
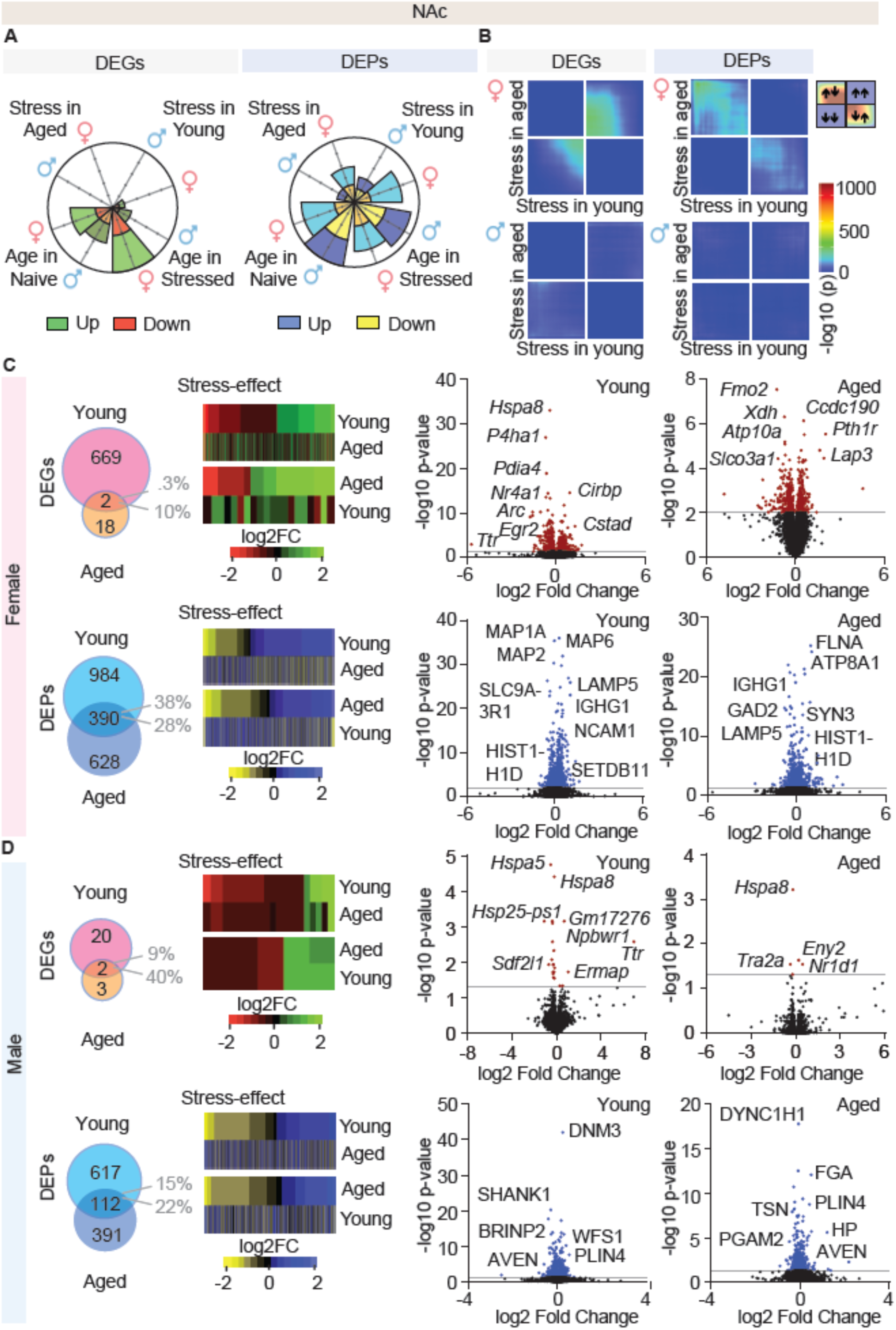
In the NAc, there is little concordance of stress effects across age. **A**) Petal plots summarize the numbers of up- and downregulated DEGs and DEPs. **B**) RRHO plots indicate minimal association between stress effects in young versus aged mice across cohorts. **C**, **D**) Venn diagrams and heatmaps illustrate the magnitude and direction of overlapping DEGs and DEPs that are regulated by stress across age-groups. Heatmaps display significant stress-effects in one age group (first row), alongside with the fold-changes of the identical set within the second age group. They indicate a modest to no association of stress effects across age groups. Volcano plots depict top altered molecules by log10 p-value. Labels are selected for most strongly altered and representative gene products. **C**) Females. **D**) Males.

Qualitatively, stress in young mice induced classic depression-associated immediate early gene programs (e.g. *Egr2*, *Arc*, *Fos*, *Fosb*), whereas aged mice engaged a unique transcriptional profile including resilience-associated chromatin and synaptic regulators, such as *Zfp366* (PFC and NAc) and *Kdm6b* (PFC) ^22,23^.

Proteomic alterations further supported distinct age-dependent molecular reprogramming of the stress response, with a partial inverse regulation of stress-linked chromatin regulators (SIRT2, HIST1H4A and HIST1H1D) in young versus aged cohorts (**Fig. 2**, **3, Fig. S3**). Taken together, these data demonstrate that chronic stress activates distinct molecular programs in young versus aged cohorts.

### Stress is associated with aging young mice, but opposes aging in old cohorts

To test, whether stress engages aging-associated molecular programs, we overlaid stress-regulated DEG and DEP changes with aging-associated signatures derived from naïve young and aged cohorts. Aging itself induced known markers including GFAP, C4B, DPYSL3 and TUBB2B ^24–26^. Main age-regulated pathways included synaptic signaling, amino acid, translation and lipid metabolism and were in part sex-specific (**Fig. S4**).

Next, we compared stress-effects in aged mice with aging. Comparable proportions of DEGs and DEPs were regulated by both, stress in aged mice, and aging, compared to overlap between stress in young mice and aging (**Fig. 4A**). Interestingly, heatmaps show that the direction of regulation was opposite across conditions: Gene products that were upregulated by aging were downregulated by stress and vice versa (**Fig. 4B, Fig. S3**). These data were consistent across gene products, brain regions and sexes and present correlative evidence that pathways of aging may be regulated in an opposite direction by stress in aged mice. RRHO analyses and Pearson correlations further demonstrated strong positive associations between stress effects in young mice and aging across sexes and brain regions (**Fig. 4C**, **Table S2**).

**Fig. 4:**
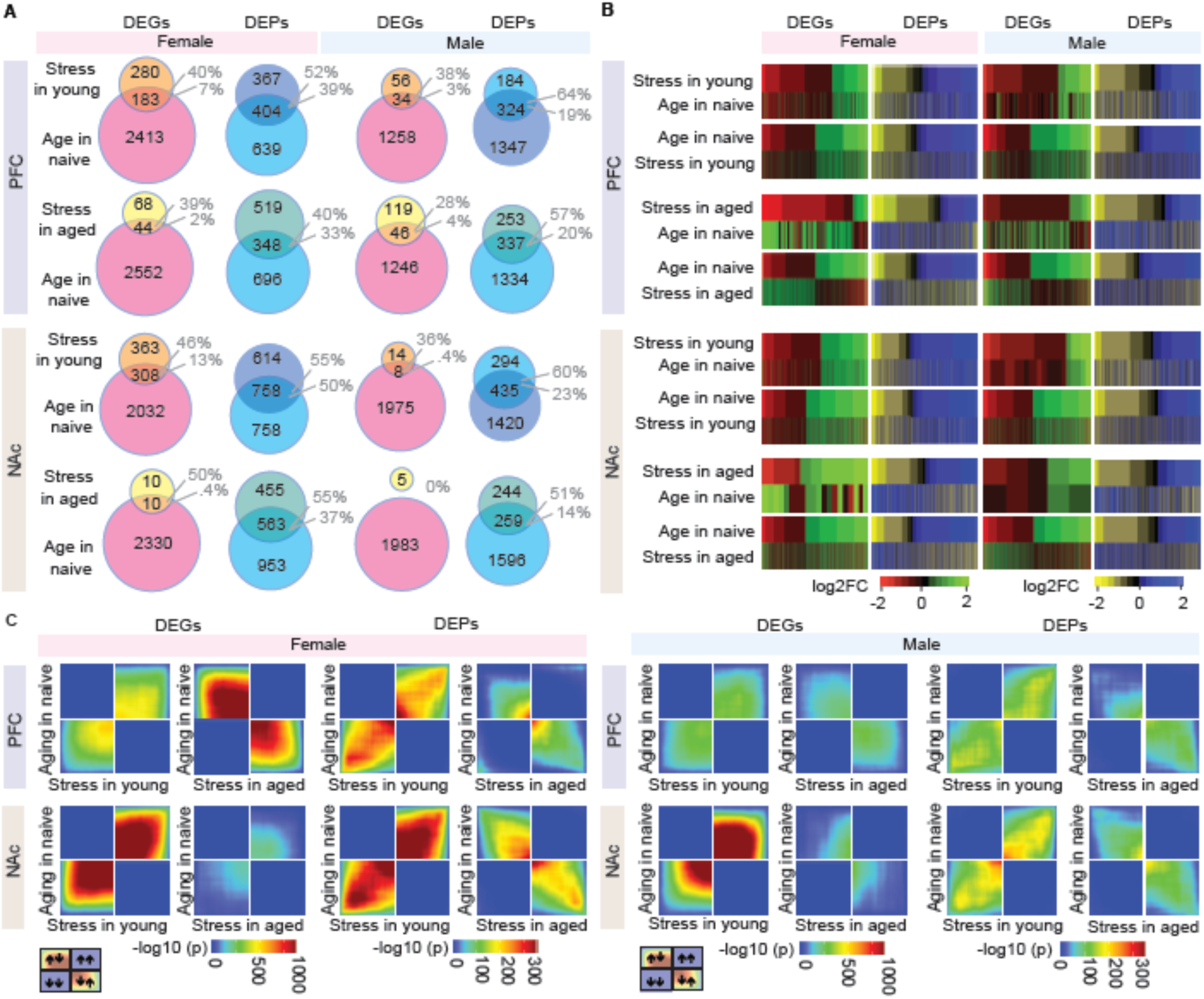
Co-regulation of gene products in aging and young, stressed mice but inverse regulation in aging and aged, stressed mice. **A**) Venn diagrams show that the overlap between age- and stress effects is comparable for stress-effects in young and aged cohorts across brain regions, gene products and sexes. **B**) Heatmaps and **C**) RRHO plots show that across brain regions, sexes and gene products there is a positive association between aging and stress in young mice, which is inverted between stress conditions in aged mice.

In contrast, stress responses in aged mice exhibited a widespread inverse concordance with aging. Although a substantial proportion of DEGs and DEPs overlapped between stress in aged mice and aging (**Fig. 4A**), the directionality was systemically reversed: DEGs upregulated during aging were mostly downregulated by stress and vice versa (**Fig. 4B, Fig. S3**, **Table S2**). RRHO plots indicate strong negative association between stress effects in aged mice and aging across sexes and brain regions (**Fig. 4C**).

Metascape pathway enrichment analysis established distinct stress-induced pathways in young and aged mice. In young mice, stress-regulated networks were enriched for growth factor signaling, circadian rhytms (PFC) and synapse organization (NAc) consistent with previous literature ^17,27^. They also support for a decoupling of transcriptional and translational processes by stress in both brain regions (**Fig. S5**). In aged mice, stress pathways differ from those of young mice and point towards tissue remodeling and angiogenesis changes in the PFC as previously unknown mechanisms in the stress response.

In summary, stress has a distinct, age-specific component. Whereas stress in young mice partially converges on aging-associated programs in young mice (“accelerated aging”), stress in aged mice inversely aligns with age-effects at a comparable scale (“rejuvenation”).

### Aging amplifies sex-differences in the brain’s stress response

To determine the interplay between sex, aging, and chronic stress, we compared associated molecular changes.

In agreement with previous literature ^5^, stress in young mice only showed a partial concordance between sexes. Across brain regions, a fraction of stress-regulated DEGs and DEPs overlapped between males and females (**Fig. 5A**). Heatmaps showed a moderate association across sexes in both brain regions, which was weaker for DEPs (**Fig. 5B**,).

**Fig. 5:**
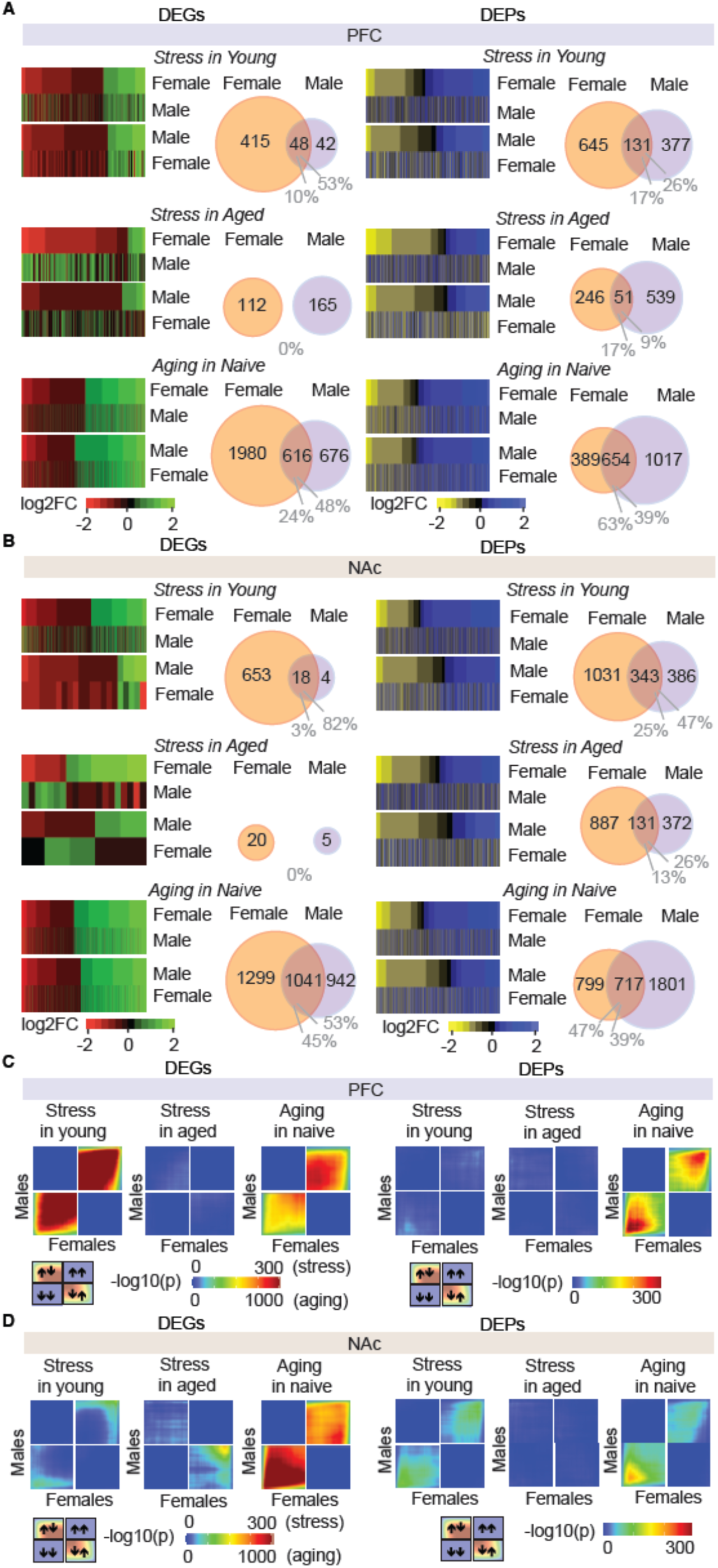
Sex-specificity of stress effects becomes more pronounced with age. **A**) PFC. **B**) NAc. **A**, **B**) Stress- and age-effects were compared between sexes using heatmaps and Venn diagrams. On the RNA-level, allover fewer changes were detected in males. Heatmaps depicted a modest association of stress-effects in young mice across sexes, further reflected by a modest overlap in Venn diagrams (DEGs and DEPs). In aged mice, the overlap in stress-effects between sexes was diminished on both, RNA and protein levels. Aging effects showed a comparatively large overlap across sexes as well an association in heatmaps. **C, D**) RRHO plots further confirm that aging increases the sex-specificity in the stress-response. Aging shows a profound association across sexes in both brain regions and measures (note the different scales for stress and aging). **C**) PFC. **D**) NAc.

Interestingly, in aged mice, stress responses became strongly sex-specific. No stress-regulated DEGs and only a small proportion of DEPs overlapped between males and females, suggesting that the stress responses become more sex-specific with age (**Fig. 5A**). Accordingly, heatmaps revealed weak to inverse associations between sexes in aged mice after stress (**Fig. 5A, B**). These data indicate that age has an impact on sex-specific molecular response to stress.

In contrast, aging-associated differential results remained comparatively conserved between sexes. Across brain regions, a higher proportion of age-regulated DEGs and DEPs, overlapped between sexes, compared to stress (**Fig. 5A**, lower part), with consistent evidence of positive association between sexes in heatmaps and RRHO plots (**Fig. 5B-D**).

Taken together, these results demonstrate that the stress response becomes more sex-specific during aging. This identifies sex as an increasingly dominant biological variable in stress biology with advancing age.

### Aging-dependent inversion of stress programs extends to the periphery

To assess whether the age-dependent inversion of stress programs is restricted to the brain, we performed quantitative proteomics on liver tissue from young and aged mice following CVS (**Fig. 6**).

**Fig. 6:**
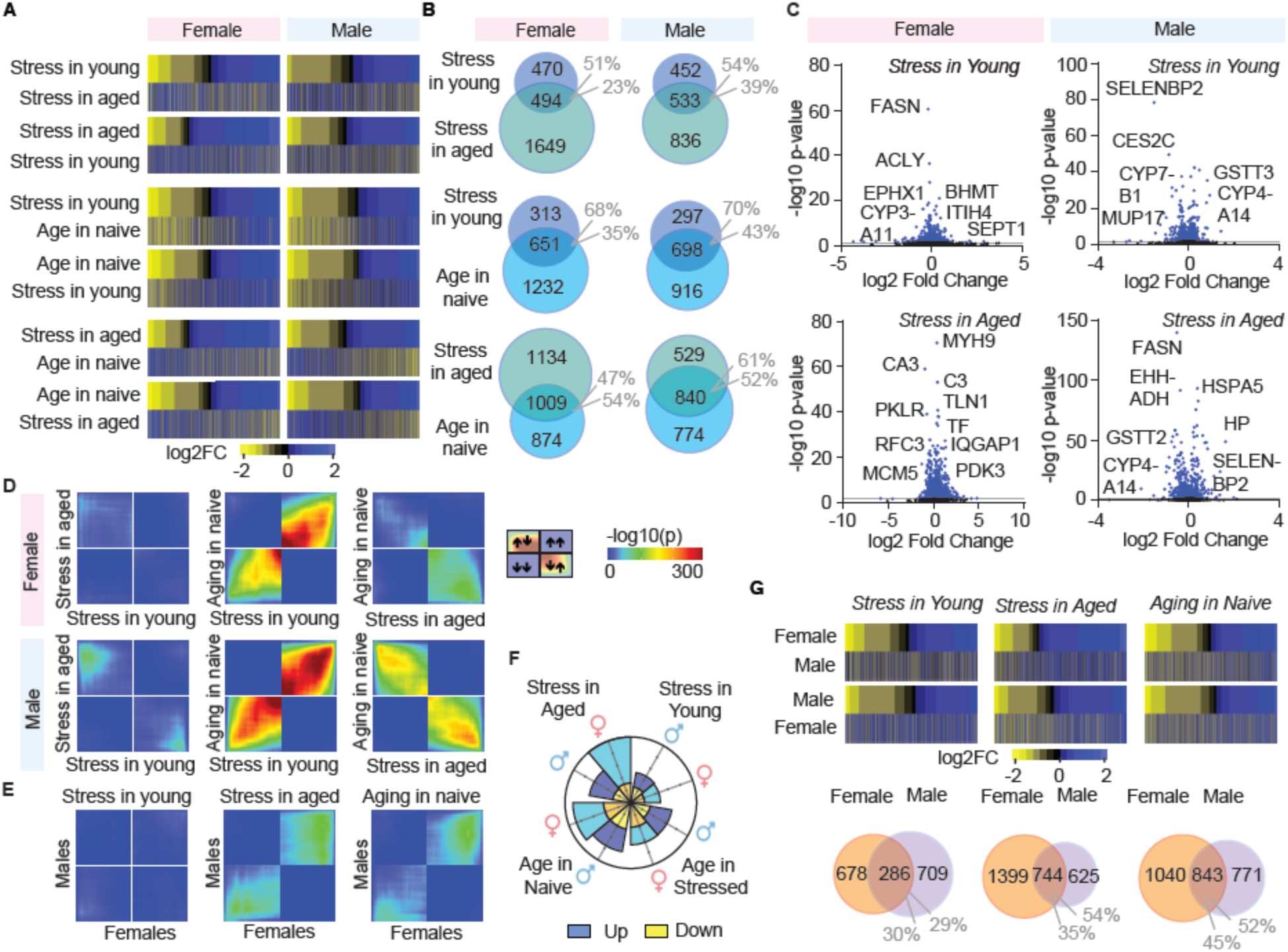
Proteomics on the liver shows an age-dependent stress effect. **A**) Heatmaps show no association of stress-effects across ages, but a modest association between stress in young mice and aging, as well as a modest reverse association between stress in aged mice and aging. **B**) DEPs in Venn diagrams similarly overlap between stress effects and aging across age cohorts. **C**) Volcano plots across stress conditions highlight selected DEPs with partially inverse regulation during aging (e.g. SELENBP2 and CYP4A14 in males). **D**) In both sexes, the stress response between young and aged mice shows little evidence of similarity in RRHO plots (left). Stress in young mice and aging are positively associated, while stress in aged mice is not. **E**) In the liver, sex-differences in the stress response are most pronounced in young mice and become more similar with age, comparable in scale to sex-differences in aging. **F**) Petal plots depict total numbers of DEPs and their direction of regulation. **G**) Heatmaps show no association across sexes in the stress response of young mice with a modest overlap shown in Venn diagrams. Stress in aged mice and aging show a moderate overlap and a moderate positive association across sexes. Volcano plots depict top altered molecules by log10 p-value. Labels are selected for most strongly altered and representative gene products.

As in the brain, stress evoked age-specific and sometimes opposing protein remodeling in the liver. Stress-regulated protein patterns showed a weak concordance between young and aged cohorts. However, similar to the brain, stress in young mice is associated with aging, whereas stress in aged mice shows an inverse pattern. This was reflected by substantial overlaps in significantly regulated gene products between stress and aging across age groups (**Fig. 6A-C**, **Fig. S3**).

Accordingly, RRHO plots depict patterns similar to the brain, with no association of stress effects in young versus aged mice, a positive association between stress in young mice and aging, and a negative association between stress in aged mice and aging (**Fig. 6D**). Effect sizes of the stress response during aging are comparable across age cohorts, albeit with an inverse direction. These data establish that the age-dependent inversion of stress responses during aging is systemic and not confined to the brain.

Sex-stratified analyses identified a distinct pattern in the liver compared to the brain. Livers of young mice showed strong sex-specific stress responses, whereas they became more sex-conserved in aged mice and approached the degree of sex conservation observed for aging-associated programs. Thus, unlike the brain, where sex specificity of stress responses increases with age, the liver exhibits a convergence of stress programs across sexes with advancing age (**Fig. 6E-G**).

### Stress remodels the aged prefrontal brain vasculature in an anti-aging direction

To identify candidate mechanisms underlying the age-dependent inversion of stress effects, we focused on the angiogenesis-associated gene network, which emerged as the most stress-regulated pathway cluster in the aged female PFC. Endothelial bleeding, blood vessel integrity, and blood-brain barrier (BBB) are altered by chronic stress ^28–30^ and aging ^31–33^. However, how brain vasculature is affected by the interplay of stress and aging is currently unknown.

Consistent with the stress-aging inversion observed globally, angiogenesis-related DEGs in aged mice were inversely regulated compared to aging (**Fig. 7A**). Analysis of corresponding DEPs identified a similar pattern (**Fig. 7B**). Notably, four out of the five gene products that showed overlap between stress-regulated DEGs and DEPs within aged females (*Fn1*, *Gpc4*, *Iqgap* and *Vwf*), belonged to the angiogenesis network, indicating that vascular remodeling is a focal point of age-dependent stress programs.

**Fig. 7:**
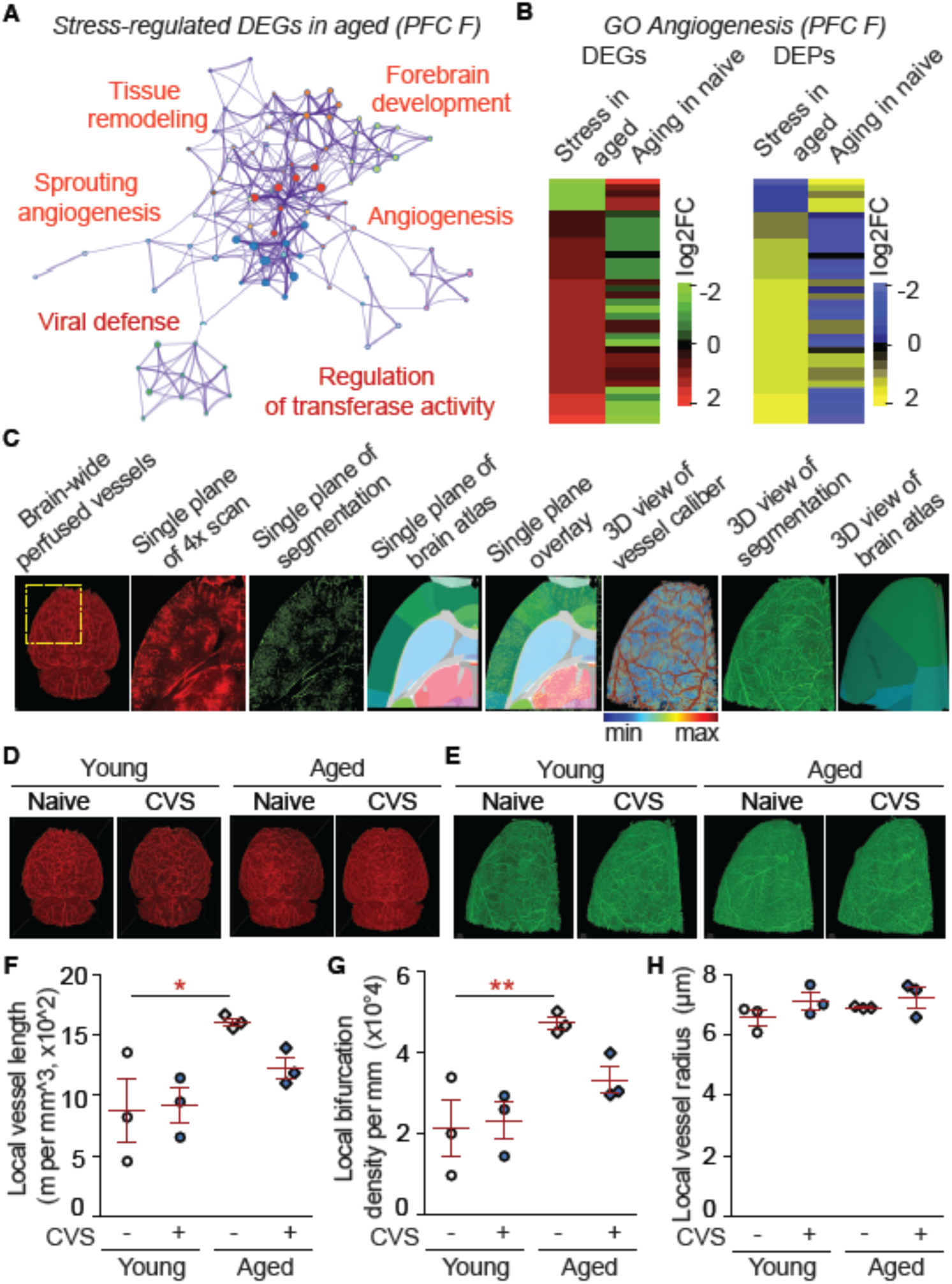
Angiogenesis-associated gene products are altered by stress and aging (females). **A**-**C**): DEGs. Green: up-regulated, red: down-regulated. **A**) Metascape clusters of stress-regulated genes in aged mice. **B**) Heatmaps of angiogenesis cluster. Left: DEGs, Right: DEPs corresponding to cluster in Color coding: bright green: signif. up-regulated, dark green: n.s. up-regulated, bright red: signif. down-regulated; dark red: n.s. down-regulated. **C**) VesSAP segmentation work flow of a representative sample. Single planes are from 1950 µm depth. **D**) Brain-wide perfused vasculature of representative samples. **E**) VesSAP CNN segmentation of samples from **D**). **F**-**H**) Vessel imaging at the frontal pole. Inserts depict the assessed parameters in green. Statistics: 2-way ANOVA & Bonferroni *post hoc* test. Independent data points are plotted and means **±** s.e.m. are shown. n=3 per group. **F**) The local vessel length is increased by aging in naïve mice. Age effect: F(1,8) = 11.18, *P < 0.05; *post hoc* test: age effect within naïve: **P < 0.01, within CVS: n.s. **G**) The local bifurcation density is increased by aging in naïve mice. Age effect: F(1,8) = 15.82, **P < 0.01; *post hoc* test: age effect within naïve: **P < 0.01, within CVS: n.s. **H**) The local vessel radius is not affected by stress or aging. All comparisons: n.s

To determine whether these molecular signatures translate into structural vascular remodeling, we performed three-dimensional imaging on the frontal brain of female mice as previously described ^34^ (**Fig. 6C-E**). Consistent with our Omics data, in the frontal pole of the brain, aged naïve but not aged stressed mice, exhibited increased vessel local length and branching, while the vessel radius was not affected (**Fig. 7F-H**). Other frontal cortical regions showed a similar trend, albeit with more regional variations (**Fig. S6**). Taken together, these data indicate that chronic stress counteracts age-associated angiogenic remodeling in the PFC, providing a mechanistic substrate for anti-aging-like molecular and synaptic programs engaged in the aged brain.

## Discussion

We observed that young and aged mice show fundamentally distinct stress responses. Chronic stress in young mice engages programs in line with accelerated aging, while in aged mice, stress opposes aging programs at a comparable scale. This inversion extends from brain regions to the periphery and stress engages specific angiogenic, synaptic and molecular networks. These data establish age as a key biological variable in the stress response.

The accelerated aging hypothesis has been supported by mounting evidence from young and midlife cohorts, but our findings demonstrate that this framework is incomplete. While stress and aging converge in the young brain, they diverge in the aged brain. Thus, stress cannot be universally conceptualized as an aging accelerator, but instead engages age-specific molecular programs whose directionality reverses in late life.

This age-dependent inversion has direct implications for psychiatric modeling and drug discovery. Most antidepressant targets are explored in young-animal stress models. While the CVS model is generally well suited for this undertaking ^5,16^, our findings suggest that these models may systematically misrepresent stress biology in late life, potentially explaining the limited efficacy of many antidepressants in elderly populations and encouraging age-stratified target discovery.

The time course of chronic stress responses across ages should be explored in more detail. An incidental study exploring one week of mild stress in a setting similar to ours (two and 24 months old C57Bl/6 mice) observed that hippocampal learning and *Bdnf* levels are impaired in young stressed, but inversed in aged stressed mice, suggesting that even at mild or early stages of stress induction, an age diversion can take place ^35^.

Aging selectively amplifies sex specificity in neural stress programs while preserving sex-conserved aging trajectories. In contrast, in the liver, stress programs converge between sexes during aging. This suggests that aging promotes sex-specific neural adaptation strategies layered upon a shared metabolic stress architecture. Underlying programs may be mediated by well-known changes in sex hormones during aging, and experience of child rearing. How these selectively affect neural versus peripheral molecular programs, may be a topic of future investigation.

Our Omics analyses show distinct regulatory mechanisms after stress and aging on the levels of RNA and proteins. While DEGs are mostly downregulated by stress, this effect is not translated to the protein level, suggesting a decoupling of RNA and proteins after stress. Candidate mechanisms include chromatin remodeling (including altered HIST1D1), altered mRNA processing, as well protein folding, catabolism, complex assembly and amino acid metabolism (see metascapes), pointing to a multi-regulatory framework of stress responses in the aged brain that deserves further investigation.

Age-dependent reprograming of stress responses may reflect distinct adaptive strategies. Energy-conserving stress programs in young animals may favor future reproductive investment, whereas active remodeling programs in aged mice may promote maintenance and protection of existing offspring or social groups. Whether similar adaptive reprograming takes place in humans remains an important open question.

Together, these findings establish aging as a reprogrammer of stress biology and motivate age-stratified approaches to modeling, mechanistic discovery and therapeutic development in stress-associated conditions.

## Acknowledgments

This project was made possible by generous funding from the Carl-Zeiss-Foundation (IMPULS #P2019-01-0006). We thank the Core Facility “Next Generation Sequencing” headed by Dr. Marco Groth for help performing RNA-sequencing. The FLI is a member of the Leibniz Association and is financially supported by the Federal Government of Germany and the State of Thuringia. This work was supported by the Vascular Dementia Research Foundation, Deutsche Forschungsgemeinschaft (DFG, German Research Foundation) under Germany’s Excellence Strategy within the framework of the Munich Cluster for Systems Neurology (EXC 2145 SyNergy, ID 390857198), ERC Consolidator Grant (A.E., GA 865323), Nomis Heart Atlas Project Grant (A.E., Nomis Foundation). Some of the graphical illustrations used in the manuscript were prepared using BioRender.com.

## Contributions

G.S., L.L., O.E., I.H., and M.K. performed mouse experiments and analysis. G.S., O.E., L.L., and M.K. planned experiments and handled paperwork involving animal experiments. G.S. and L.L. double-checked data and statistics. L.L. and I.H. analyzed dendritic spines. K.R. and S.H. analyzed the RNA-sequencing raw data, G.S. and O.E. analyzed the transcriptomics and proteomics annotated data. E.C. and N.P. obtained and analyzed proteomics raw data. C.A.H. and K.L.R. provided infrastructure or mice, and conceptual advice and edited the manuscript. M.I.T., F.H. and A.E. performed 3D brain vasculature measurements and analysis. O.E. designed and planned the study, obtained funding and wrote the article.

## Competing interests

A.E. is a co-founder of Deep Piction. The other authors declare no competing interests.

## Notes

https://massive.ucsd.edu/ProteoSAFe/private-dataset.jsp?task=660033d2b688454a8faed0b423935180

https://www.ncbi.nlm.nih.gov/geo/query/acc.cgi?acc=GSE317903

