## Supplemental information for "Aging reprograms the response to chronic stress"

### Supplemental material

**Statistics for figure legend 1.**

**B**) Forced swim test. n = 21,18,18,18; 2-way ANOVA: Interaction stress x age: $F(1,71) = 29.23$, $P < 0.0001$; Bonferroni *post hoc* test: stress effect within young: ***$P < 0.001$, within aged: **$P < 0.01$; age-effect within naïve: **$P < 0.01$, within CVS: ***$P < 0.001$.

**C**) Splash test. n = 11,12,13,13; 2-way ANOVA: Interaction stress x age: $F(1,45) = 13.65$, $P < 0.001$; Bonferroni *post hoc* test: stress effect within young: **$P < 0.01$; age-effect within CVS: ***$P < 0.001$.

**D**) Body weight. n = 40,40,36,37; 2-way ANOVA: stress effect: $F(1,149) = 181.70$, $P < 0.0001$; age effect: $F(1,149) = 57.84$, $P < 0.0001$; Bonferroni *post hoc* test: stress effect within young: ***$P < 0.001$, within within aged: ***$P < 0.001$; age effect within naïve: **$P < 0.01$, within CVS: ***$P < 0.001$.

**G**) Neck-containing spine density. n = 76,75,77,78 dendrites from 8 mice; 2-way ANOVA: Interaction stress x age: $F(1,302) = 63.92$, $P < 0.0001$; Bonferroni *post hoc* test: stress effect within young: ***$P < 0.001$; within aged: ***$P < 0.001$; age effect within naïve: ***$P < 0.001$; within CVS: ***$P < 0.001$.

**H**) Cumulative spine head diameter. Gehan-Breslow-Wilcoxon Test: Stress effect within young: $\chi^2 = 1.09$, df = 1, *$P < 0.05$, within aged: $\chi^2 = 9.93$, df = 1, **$P < 0.01$; age effect within naive: $\chi^2 = 44.81$, df = 1, ***$P < 0.0001$, within CVS: $\chi^2 = 1.60$, df = 1, $P > 0.05$. For better visuability, the x-axis arbitrarily set to 0-6 µm. Spines with a larger diameter were however included in the analysis.

**J**) Body temperature in young mice. Head. n = 15,16,15,15,17,18,18,18; 2-way ANOVA: stress effect: $F(1,124) = 12.76$, $^{\#\#\#}P < 0.0001$. Tail. n = 15,15,16,15,17,17,18,18; stress effect: $F(1,123) = 4.34$, $^{\#}P < 0.05$. Bonferroni *post hoc* test: n.s.

**K**) Body temperature in aged mice. Head. n = 17,18,17,17,17,17,17,17; 2-way ANOVA: all comparisons: n.s. Tail. n = 16,17,18,17,16,16,16,17; stress effect: $F(1,125) = 4.70$, $^{\#}P < 0.05$. Bonferroni *post hoc* test: n.s.

**L**). Body temperature after stress induction on day 21. n = 16,17,17,16,16,17; 2-way ANOVA: stress effect: $F(1,88) = 21.00$, $^{\#\#\#}P < 0.0001$; factor of body part: $F(2,93) = 3.58$, $P < 0.05$; Bonferroni *post hoc* test: age effect within tail: **$P < 0.01$.

### Table S1: Metrics on altered gene products.

| | | Female | | | Male | | |
|---|---|---|---|---|---|---|---|
| Region | Parameter | Stress in young | Stress in aged | Aging in naive | Stress in young | Stress in aged | Aging in naive |
| Transcriptomics PFC | total sign. (no NAN/Krt) | 463 | 112 | 2597 | 90 | 165 | 1292 |
| | % up | 34.13 | 14.29 | 48.48 | 30.00 | 18.79 | 56.66 |
| | Total detected DEGs (no Padj "NA") | 14,895 | 17,029 | 17,741 | 13,475 | 9,200 | 18,455 |
| | % sign. changed | 3.11 | 0.66 | 14.64 | 0.67 | 1.79 | 7.00 |
| Transcriptomics NAc | total sign. (no NAN/Krt) | 671 | 20 | 2340 | 22 | 5 | 1984 |
| | % up | 44.71 | 0.65 | 58.89 | 22.73 | 40.00 | 53.18 |
| | Total detected DEGs (no Padj "NA") | 18,523 | 16,375 | 22,100 | 17,807 | 15,658 | 19,954 |
| | % sign. changed | 3.62 | 0.12 | 10.59 | 0.12 | 0.03 | 9.94 |
| Proteomics PFC | total sign. (no NAN/Krt) | 784 | 865 | 1043 | 508 | 590 | 1671 |
| | % up | 55.23 | 36.76 | 59.35 | 45.47 | 35.93 | 58.29 |
| | Total detected DEPs | 5,444 | 5,455 | 5,445 | 5,283 | 5,286 | 5,285 |
| | % sign. changed | 14.40 | 15.86 | 19.16 | 9.62 | 11.16 | 31.62 |
| Proteomics NAc | total sign. (no NAN/Krt) | 1374 | 1018 | 1516 | 729 | 503 | 1855 |
| | % up | 65.29 | 50.29 | 62.20 | 45.68 | 44.14 | 40.75 |
| | Total detected DEPs | 5,464 | 5,472 | 5,463 | 6,090 | 6,082 | 6,086 |
| | % sign. changed | 25.15 | 18.60 | 27.75 | 11.97 | 8.27 | 30.48 |
| Proteomics Liver | total sign. (no NAN/Krt) | 964 | 2144 | 1884 | 995 | 1369 | 1614 |
| | % up | 52.90 | 70.01 | 52.92 | 47.44 | 56.32 | 46.03 |
| | Total detected DEPs | 4,974 | 5,009 | 4,990 | 4,189 | 4,191 | 4,188 |
| | % sign. changed | 19.38 | 42.80 | 37.76 | 23.75 | 32.67 | 38.54 |

### Table S2: Pearson correlations.

| | PFC | | | | NAc | | | | Liver | |
|---|---|---|---|---|---|---|---|---|---|---|
| | DEG | DEP | DEG | DEP | DEG | DEP | DEG | DEP | DEP | DEP |
| | F | F | M | M | F | F | M | M | F | M |
| Stress in young vs aging in naive | 0.93 | 0.81 | 0.97 | 0.74 | 0.93 | 0.64 | NA | 0.66 | 0.6 | 0.78 |
| Stress in aged vs aging in naive | -0.94 | -0.23 | -0.53 | -0.49 | -0.83 | -0.62 | NA | -0.56 | -0.33 | -0.58 |
| Stress in young vs stress in aged | NA | 0.56 | 0.94 | 0.04 | NA | -0.49 | NA | -0.1 | -0.09 | -0.22 |

### Tables S3-S12: Excel files on Omics data.

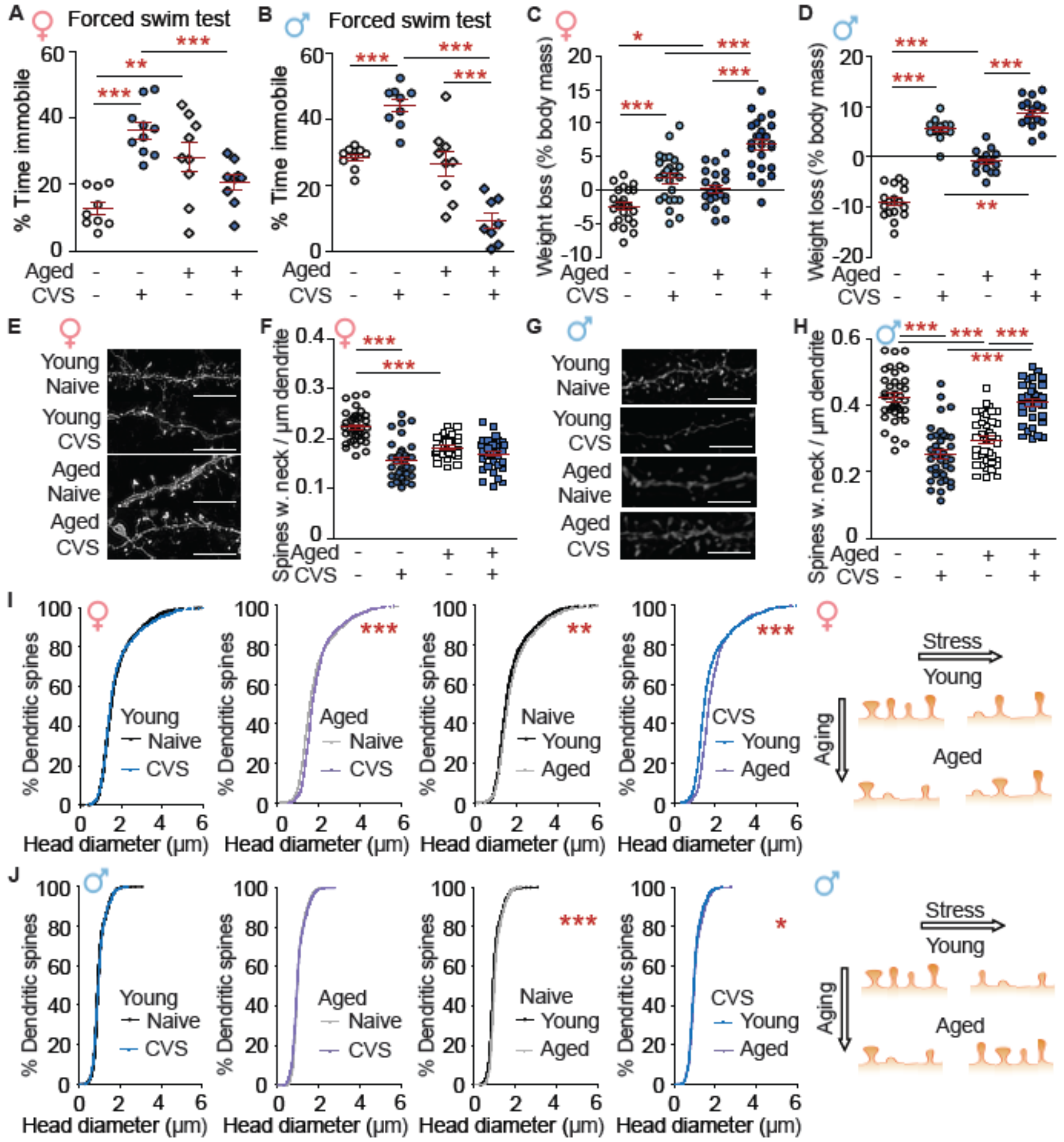


**Fig. S1: The stress by aging interaction has slight sex-specific differences. A**-**D**, **H**) Two-way ANOVA & Bonferroni *post hoc* test. **A**, **B**) Forced swim test. **A**) Females. Aged mice show no stress-induced immobility time changes but generally have an increased immobility time. n = 9-10; Interaction stress x age: $F(1,33) = 29.23$, $P < 0.001$; *post hoc* test: stress effect within young: ***$P < 0.001$; age effect within naïve: **$P < 0.01$, within CVS: ***$P < 0.001$. **B**) Males. Stress increases immobility time in young but reduces immobility time in aged mice. n = 8-9; Interaction stress x age: $F(1,31) = 44.99$, $P < 0.0001$; *post hoc* test: stress effect within young: ***$P < 0.001$, within aged: ***$P < 0.001$; age effect within CVS: ***$P < 0.001$. **C**, **D**) Body weights are reduced by CVS in both sexes and ages. **C**) Females. n = 22-24; stress effect: $F(1,89)$ =

59.73, P < 0.001; age effect: F(1,89) = 29.98, P < 0.001; *post hoc* test: stress effect within young: ***P < 0.001, within aged: ***P < 0.001; age effect within naïve: **P < 0.01, within CVS: ***P < 0.001. **D**) Males. n = 16-17; interaction stress by aging: F(1,62) = 14.81, P < 0.001; *post hoc* test: stress effect within young: ***P < 0.001, within aged: ***P < 0.001; age effect within naïve: ***P < 0.001, within CVS: **P < 0.01. **E**-**J**) Dendritic spine morphology. **E**, **F**, **I**) Females. **G**, **H**, **J**) Males. **E**) Females: Representative dendrites. Scale bar 10 µm. **F**) CVS reduces neck-containing spines in young females. n = 35-37 dendrites from 4 mice per group; Interaction stress by age: F(1,142) = 30.68, P < 0.0001; *post hoc* test: stress effect within young: ***P < 0.001, within aged: n.s.; age effect within naïve: ***P < 0.001; within CVS: n.s. **G**) Males: Representative dendrites. Scale bar 10 µm. **H**) CVS reduces the density of neck-containing spines in young females and increases it in aged mice. n = 37-39 dendrites from 4 mice per group; Interaction stress by age: F(1,148) = 147.20, P < 0.0001; *post hoc* test: stress effect within young: ***P < 0.001, within aged: ***P < 0.001; age effect within naïve: ***P < 0.0001, within CVS: ***P < 0.001. **I**) In females, the cumulative head diameter is increased by CVS in aged mice and by aging. Gehan-Breslow-Wilcoxon Test: Stress-effect within young: $\chi^2$ = 0.40, df = 1, n.s.; Stress-effect within aged: $\chi^2$ = 26.42, df = 1, ***P < 0.0001; age effect within naive: $\chi^2$ = 9.36, df = 1, **P < 0.0001, within CVS: $\chi^2$ = 3.89, df = 1, *P < 0.05. **J**) In males, the cumulative head diameter is not altered by CVS in young mice but increased by aging. Gehan-Breslow-Wilcoxon Test: Stress effect within young: $\chi^2$ = 0.17, df = 1, n.s., within aged: $\chi^2$ = 2.32, df = 1, n.s.; age effect within naive: $\chi^2$ = 18.66, df = 1, **P < 0.01,within CVS: $\chi^2$ = 1.60, df = 1, P > 0.05. **A**-**D**, **F**, **H**) Independent data points are plotted and means **±** s.e.m. are shown. **I**, **J**) For better visibility, the x-axis arbitrarily set to 0-6 µm. Spines with a larger diameter were however included in the analysis.

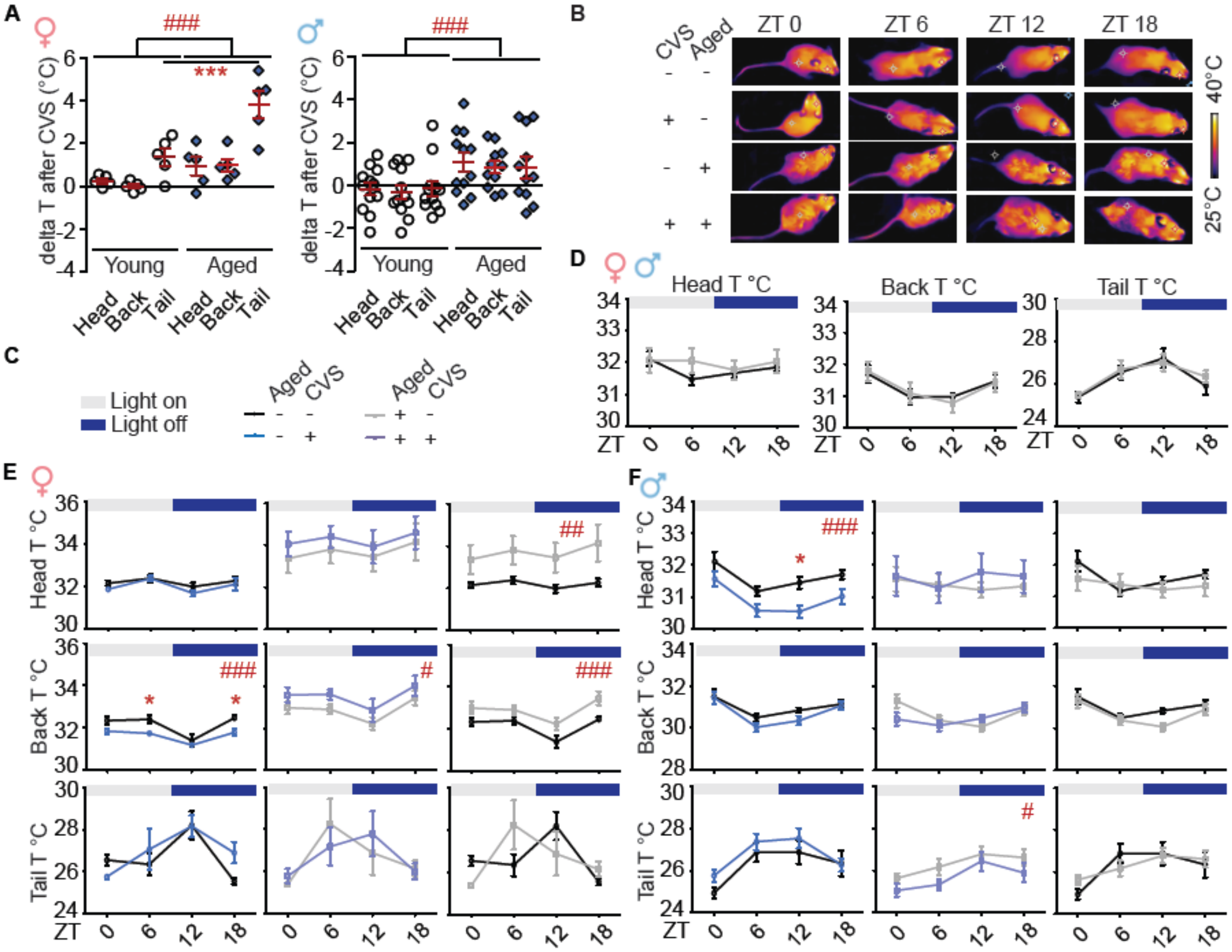


**Fig. S2: Stress-induced body temperature changes have slight sex differences.** Statistics: 2-way ANOVA & Bonferroni *post hoc* test. **A**) The body temperature was assessed before (ZT 0) and right after CVS. The temperature was significantly more elevated after CVS in aged versus young mice of both sexes. In females, the difference was particularly evident in the tail temperature. Females: n = 5 per group; age effect: F(1,24) = 18.94, ###P < 0.0001; factor of body part: F(2,24) = 19.07, P < 0.0001; *post hoc* test: age effect within tail: ***P < 0.001. Males: n = 12,12,12,12,11,11; stress effect: F(1,64) = 13.85, ###P < 0.001. **B**) Representative images. **C**) Legend for **B-F**). **D**) Both sexes combined: No age effects on body temperature are detected. Head: n = 15,16,15,15,17,18,17,17; all comparisons: n.s. Back: n = 16,16,15,16,17,18,18,17;: age effect: F(1,125) <0.01, n.s. Tail: n = 15,15,16,15,16,17,18,17; age effect: F(1,121) = 0.21, n.s. **E**, **F**) Minor sex differences in stress- and age-dependent body temperature changes. **E**) Females. Stress reduces the back temperature in young mice and increases it in aged mice. Aged females have a higher core (head + back) temperature than young mice. Stress effect in young mice (left): Head: n = 4 (naive), 5 (CVS); stress effect: F(1,28) = 20.25, n.s. Back: n = 4 (naive), 5 (CVS); F(1,28) = 2.09, ###P = 0.0001; *post hoc* test: at ZT 6: *P < 0.05; at ZT 18: *P < 0.05. Tail: n = 4 (naive), 5 (CVS); F(1,28) = 0.59, n.s. Stress effect in aged mice (middle): Head: n = 5 per group; F(1,32) = 1.20, n.s. Back: n = 5 per group; F(1,32) = 6.32, #P < 0.05. Tail: n = 5 per group; F(1,32) < 0.01, n.s. Age effect in naïve mice (right): Head: n = 4 (young), 5 (aged);

F(1,28) = 12.25, $^{\#\#\#}P = 0.01$; Back: n = 4 (young), 5 (aged); F(1,28) = 16.04, $^{\#\#\#}P < 0.001$; Tail: n = 4 (young), 5 (aged); F(1,28) < 0.01, n.s. **F**) In young but not aged males, CVS reduces the head temperature. Stress effect in young mice (left): Head: n = 11,12,12,12,12,12,12,13: F(1,88) = 21.00, $^{\#\#\#}P < 0.0001$; *post hoc* test: ZT12: *P <0.05. Back: n = 12,12,11,12,12,12,13,13; F(1,89) = 2.64, n.s. Tail: n = 11,12,12,12,12,13,13,13; F(1,90) = 2.43, n.s. Stress effect in aged mice (middle): Head: n = 12,13,12,12,12,12,12,11; F(1,88) = 0.46, n.s. Back: n = 12,13,12,13,12,12,11,12; F(1,89) = 0.56, n.s. Tail: n = 12,12,12,13,11,11,12,12; F(1,87) = 9.11, $^{\#}P < 0.05$. Age effect in naïve mice (right): Head: n = 11,12,12,12,12,13,12,12; F(1,88) = 1.35, n.s. Back: n = 12,12,11,12,12,13,12,13; F(1,89) = 2.56, n.s. Tail: n = 11,12,12,12,12,12,12,13; F(1,88) = 0.03, n.s. ZT – Zeitgeber time. For simplicity, effects of circadian time are not depicted in the statistics. There were no interactions between circadian effects and stress or age effects detected. Only significant *post hoc* test data are described.

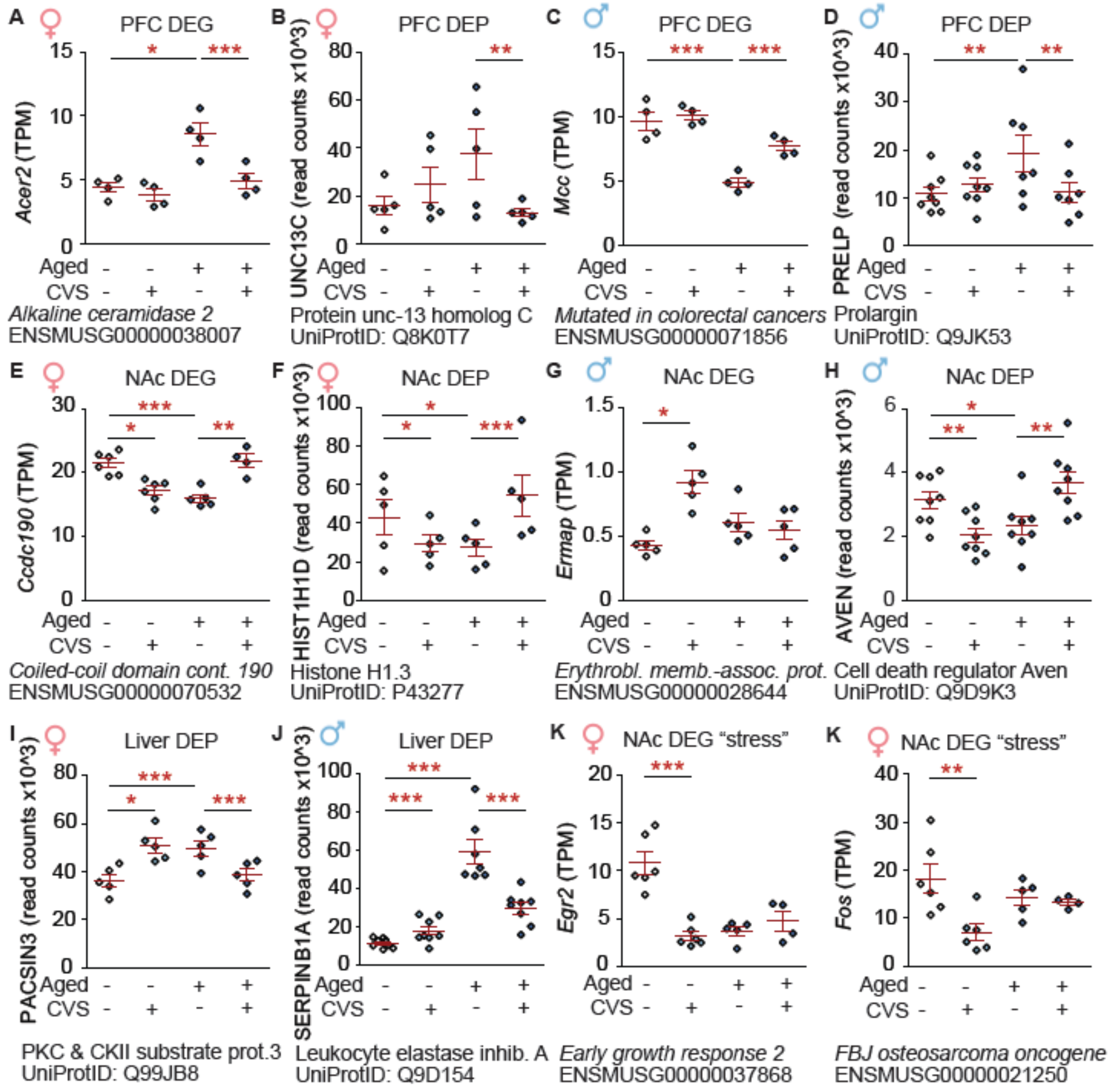

**Fig. S3: Examples of gene products that were affected by stress and aging.** Omics raw data are shown. Statistics refer to Padj from Omics analyses generated with DESeq2 and Spectronaut, respectively (*P < 0.05; ** P < 0.01, ***P < 0.001). The base mean refers to the expression across all groups. TPM = transcript per million. OE: change DEGs to cursive and DEPs to capital.

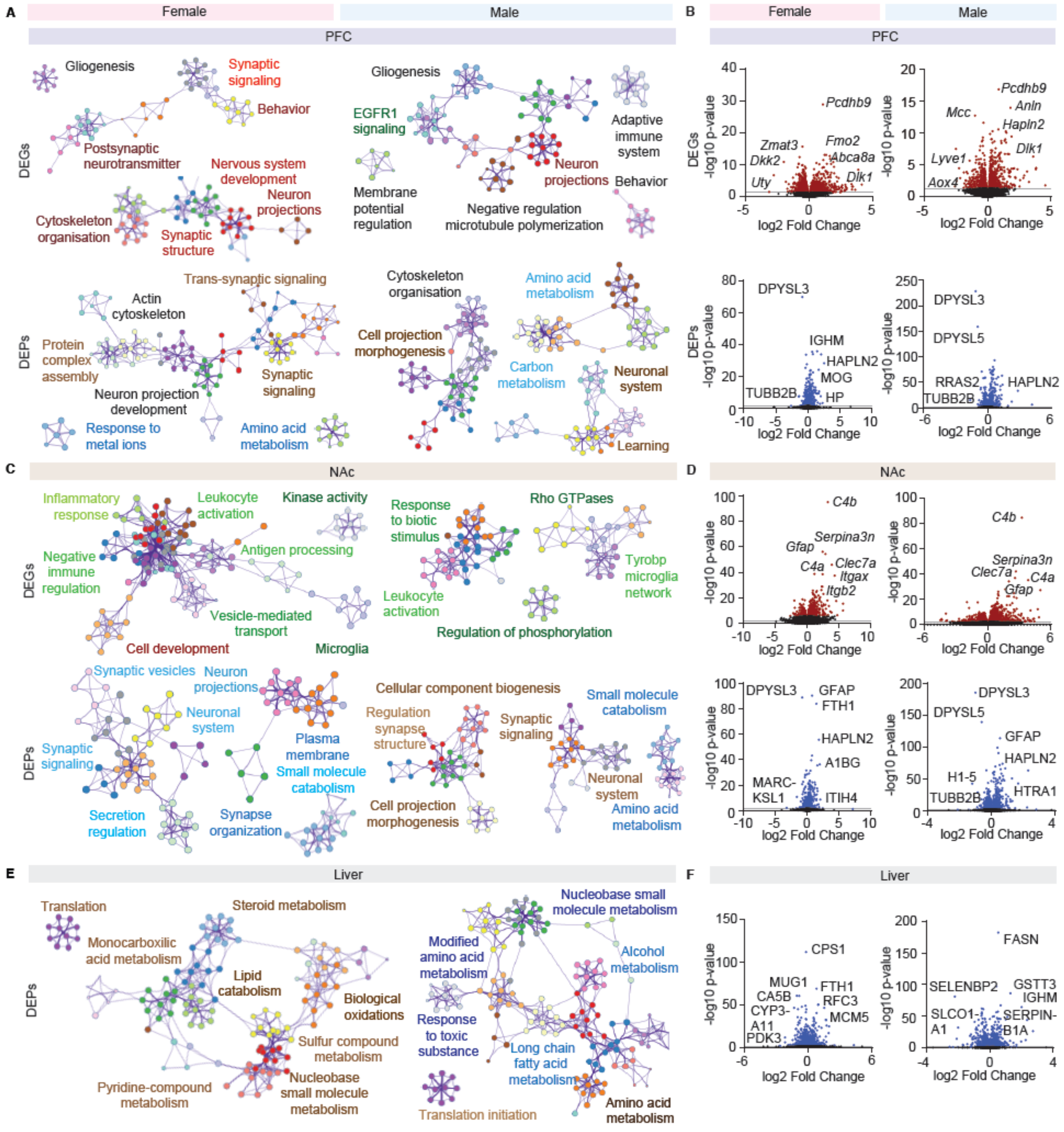


**Fig. S4: Aging effects in naïve mice. A**, **C**, **E**) Metascapes. Only the most abundant pathways are shown. DEGs: green – upregulated, red – downregulated. DEPs: blue – upregulated, yellow – downregulated. **B**, **D**, **H**) Volcano plots. **A**, **B**) PFC. **A**) Aging reduces synaptic signaling in DEGs and DEPs of females. In both sexes, behavior and learning are reduced, whereas the amino acid metabolism is increased across gene products. **B**) Volcano plots include age-related gene products such as DPYSL3 and TUBB2B. **C**, **D**) NAc. **C**) Within DEGs, there is an upregulation of immune-related processes in both sexes. Synaptic signaling may be regulated by aging in a sex-specific manner (up in females, down in males within DEPs). As in the PFC, at least in males, there is an age-related increase in the amino acid metabolism. **D**) Volcano plots show regulation of age-related DEPs such as C4A, DPYSL3 and GFAP. **E**, **F**) Liver. **E**) There are age-dependent alterations in metabolism linked to lipids (steroids, lipid catabolism, fatty acid metabolism) as well as a reduction in translation across sexes. Additionally, there may be alterations in metabolism related to small molecules, alcohol and toxic substances. **F**) This is reflected by changes in fatty acid and detoxification-related DEPs such as FASN and CYP3A11. Volcano plots depict top altered molecules by log10 p-value. Labels are selected for most strongly altered and representative gene products.

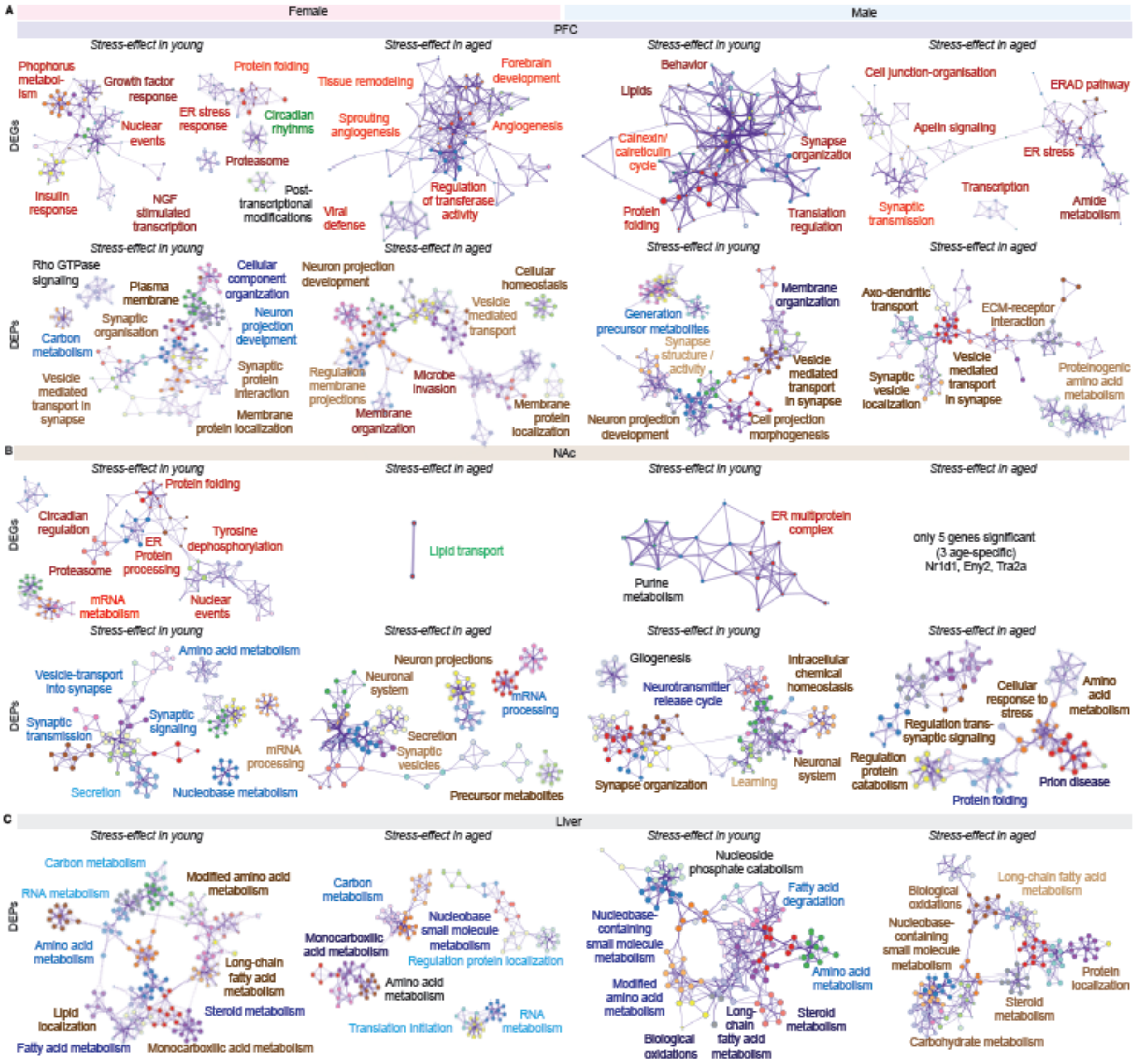


**Fig. S5: Metascapes of stress-comparisons in both age groups and sexes. A**) PFC. **B**) NAc. **C**) Liver. Similar to aging (**Fig. S4**), stress in young mice induced changes in synaptic signaling, carbon and amino acid metabolism, lipid metabolism and learning. Stress in aged mice induced some pathways that were regulated in a similar manner (e.g. carbon metabolism in the liver (**C**), some pathways that were regulated in an opposing manner (mRNA processing and synaptic vesicles in the NAc (**B**) or that were specific to aged stressed mice, such as angiogenesis and sprouting angiogenesis in PFC (**A**).

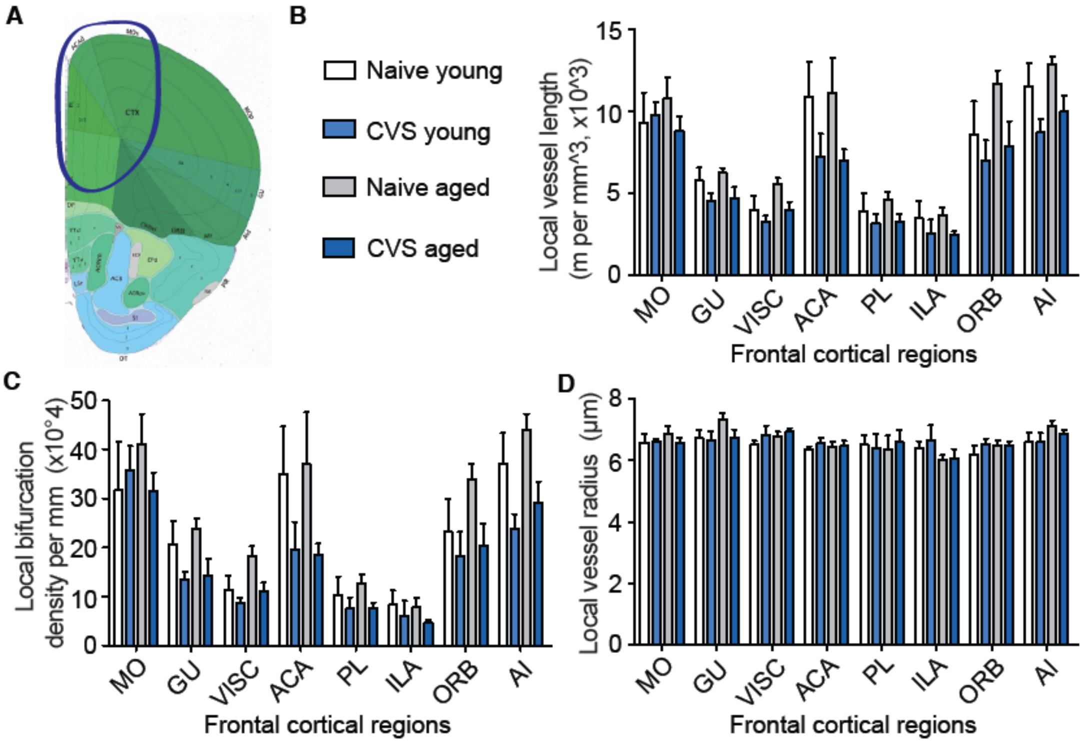


**Fig. S6: Blood vessel parameters in other frontal cortical regions. A**) Allen brain atlas coronal mouse atlas: in blue the approximate region is labeled that has been collected for Omics**.** Image represents the most posterior part collected, whereas the frontal pole represents the most anterior part. **B-D**) Frontal cortical regions other than the frontal pole. **B**) Length of blood vessels. **C**) Number of bifurcations. **D**) Vessel radius. Regions were annotated according to the Allen brain atlas: MO – somatomotor area, GU – gustatory area, VISC – visceral area, ACA – anterior cingulate area, PL – prelimbic area, ILA – infralimbic area, ORB – orbital area, AI – agranular insular area.

### Extended methods

**CVS**. A combination of the acute behavioral tests forced swim test and splash test was performed as described [1,2]. The CVS protocol was performed as described [1]. In brief, mice received 21 days of stress with one of three stressors presented in a semi-random order, where the same stressor does not occur on two consecutive days. Stressors consist of 1 h of either, tube restraint, tail suspension, or 100 mild electric random foot shocks. Behavioral experiments were conducted in the active phase (dark phase, under red light) of the light cycle, while stress induction was performed within the first half of the light phase. CVS cohorts were scored for health parameters before and after each stress induction. No adverse consequences were noted directly after the induction. The weight of mice was assessed at least once a week. Mice were housed in pairs of 2-5 unless temporary single housing necessary for behavioral testing. Forced swim and splash tests were performed as previously described [3,4]. Thermal images were acquired at four time points in a temperature-controlled room (25 ± 1 °C) using a HIKMICRO Pocket2 camera (HIKMICRO). Mean surface temperature was quantified from manually predefined regions of interest located between the ears, on the back and at the tail of the mice using HIKMICRO Analyzer software (v2.0.1.4, build 250919).

**Next-generation RNA-sequencing.** The Library was prepared using Novogene NGS RNA Library Prep Set (PT042) according to manufacturer instructions from 300ng RNA. A total of 500ng RNA input material per NAc sample was processed using NEBNext Ultra II Directional RNA Library Preparation Kit (#E7760) in combination with NEBNext Poly(A) mRNA Magnetic Isolation Module (#E7490) and NEBNext Multiplex Oligos for Illumina (Unique Dual Index UMI Adaptors RNA) (#E7416) following the manufacturer's instructions (all New England Biolabs). A final amplification of the library was performed. RNA-seq was performed by the company Novogene (U.K.) and by the sequencing core facility of the Fritz-Lipmann-Institute for Aging, Jena, Germany. An in-house RNA-sequencing analysis pipeline (https://github.com/Hoffmann-Lab/rippchen) was applied which utilized Trimmomatic [5] v0.39 (5nt sliding window, mean quality cutoff 20) for read quality trimming. According to FastQC v0.11.9 reports, the Illumina universal adapter was clipped off the 3' reads end using Cutadapt [6] v2.10. Sequencing errors were detected and corrected using Rcorrector [7] v1.0.4. Potential ribosomal RNA derived sequences were depleted by utilizing SortMeRNA [8] v2.1b. Subsequently, the data was aligned to the mouse reference genome GRCm38 (mm10) with segemehl [9,10] v0.3.4 in splice-aware mode and accuracy cutoff raised to 95%. For single-end data with unique molecular identifiers, following the extraction of unambiguously aligned reads, mappings were further deduplicated for over-amplified PCR fragments utilizing UMI-tools [11] v1.1.1. To quantify aligned fragments on Ensembl v102 reference annotation via featureCounts [12] v2.0.1 (exon-based meta-feature, minimum overlap 10nt), library strandness setting was inferred using RSeQC [13] v4.0.0. Afterwards, DESeq2 [14] v1.34.0 was applied to test for differentially expressed genes. Only significant results (Benjamini-Hochberg adjusted P-values <= 0.05) were considered in downstream analyses and interpretations.

**Proteomics.** For proteomics analysis, tissues were resuspended in PBS and lysis buffer was added to a final concentration of 5% SDS, 100 mM HEPES and 50 mM DTT. The samples were sonicated (Bioruptor Plus, Diagenode, Belgium) for 10 cycles (30 sec ON/60 sec OFF) at a high setting at 20°C, followed by boiling at 95°C for 7 min. Reduction was followed by alkylation with iodoacetamide (final concentration 15 mM) for 30 min at room temperature in the dark. Samples were acidified with phosphoric acid (final concentration 2.5%), and seven times the sample volume of S-trap binding buffer was added (100 mM TEAB, 90% methanol). Samples were bound on 96-well S-trap micro plate (Protifi) and washed three times with binding buffer. Trypsin in 50 mM TEAB pH 8.5 was added to the samples (1 µg per sample) and incubated for 1 h at 47°C. The samples were eluted in three steps with 50 mM TEAB pH 8.5, elution buffer 1 (0.2% formic acid in water) and elution buffer 2 (50% acetonitrile and 0.2% formic acid). The eluates were dried using a speed vacuum centrifuge (Eppendorf Concentrator Plus, Eppendorf AG, Germany) and they were reconstituted in in MS Buffer (5% acetonitrile, 95% Milli-Q water, with 0.1% formic acid), spiked with iRT peptides (Biognosys, Switzerland) and loaded on Evotips (Evosep) according to the manufacturer's instructions. In short, Evotips were first washed with Evosep buffer B (acetonitrile, 0.1% formic acid), conditioned with 100% isopropanol and equilibrated with Evosep buffer A. Afterwards, the samples were loaded on the Evotips and washed with Evosep buffer A. The loaded Evotips were topped up with buffer A and stored until the measurement. For LC-MS Data independent analysis (DIA), peptides were separated using the Evosep One system (Evosep, Odense, Denmark) equipped with a 15 cm x 150 µm i.d. packed with a 1.9 µm Reprosil-Pur C18 bead column (Evosep Performance, EV-1137, PepSep, Marslev, Denmark). The samples were run with a pre-programmed proprietary Evosep gradient of 44 min (30 samples per day) using water and 0.1% formic acid and solvent B acetonitrile and 0.1% formic acid as solvents. The LC was coupled to an Orbitrap Exploris 480 (Thermo Fisher Scientific, Bremen, Germany) using PepSep Sprayers and a Proxeon nanospray source. The peptides were introduced into the mass spectrometer via a PepSep Emitter 360-µm outer diameter × 20-µm inner diameter, heated at 300°C, and a spray voltage of 2 kV was applied. The injection capillary temperature was set at 300°C. The radio frequency ion funnel was set to 30%. For DIA data acquisition, full scan mass spectrometry (MS) spectra with a mass range of 350–1650 m/z were acquired in profile mode in the Orbitrap with a resolution of 120,000 FWHM. The default charge state was set to 2+, and the filling time was set at a maximum of 20 ms with a limitation of $3 \times 10^6$ ions. DIA scans were acquired with 40 mass window segments of differing widths across the MS1 mass range. Higher collisional dissociation fragmentation (normalized collision energy 30%) was applied, and MS/MS spectra were acquired with a resolution of 30,000 FWHM with a fixed first mass of 200 m/z after accumulation of $1 \times 10^6$ ions or after filling time of 45 ms (whichever occurred first). Data were acquired in profile mode. For data acquisition and processing of the raw data, Xcalibur 4.4 (Thermo) and Tune version 4.0 were used. For data processing, DIA raw data were analyzed using the directDIA pipeline in Spectronaut v.19 (Biognosys, Switzerland) with BGS settings besides the following parameters: Protein LFQ method= QUANT 2.0, Proteotypicity Filter = Only protein group specific, Major Group Quantity = Median peptide quantity, Minor Group Quantity = Median precursor quantity, Data Filtering = Qvalue, Normalizing strategy = Local Normalization. The data were searched against a species specific (*Mus musculus*, 16,747 entries, v. 160106) and a contaminants (247 entries) Swissprot database. The identifications were filtered to satisfy FDR of 1 % on peptide and protein level. Relative protein quantification was performed in Spectronaut using a pairwise t-test performed at the precursor level followed by multiple testing correction according to

Benjamini-Hochberg. The data (candidate table) and data reports (protein quantities) were then exported and further data analyses and visualization were performed with Rstudio using in-house pipelines and scripts. To select significant proteins, a q-value <0.05 were defined.

**Tissue labeling and perfusion for three-dimensional brain vasculature measurements.** Animals were anesthetized with a ketamine/xylazine mixture (administered intraperitoneally, 100 mg/kg and 16 mg/kg, respectively). After complete loss of nociceptive reflexes, the thoracic cavity was opened and 50 µl of Lycopersicon Esculentum (Tomato) Lectin (LEL, TL), DyLight 594 was injected directly into the left ventricle to circulate in the perfused vasculature for 60 s. Afterwards, the brains were extracted and post-fixed in 4% PFA for 12 h.

**Optical tissue clearing.** Brains were optically cleared using an adapted 3DISCO protocol. Briefly, we immersed them in a gradient of tetrahydrofuran (Sigma-Aldrich, 186562): 50 vol%, 70 vol%, 80 vol%, 90 vol%, 100 vol% (in distilled water), and 100 vol% at room temperature for 12 h at each concentration, delipidated in dichloromethane (Sigma-Aldrich, 270997) for 12 h at room temperature and finally incubation with the refractive index matching solution BABB (benzyl alcohol + benzyl benzoate 1:2 ratio; Sigma-Aldrich, 24122 and W213802), for at least 24 h at room temperature until transparency was achieved. Each incubation step was carried out on a laboratory shaker.

**Light-sheet fluorescence microscopy.** We used a 2× (Olympus XLFLUOR 340) and a 4x (Olympus XLFLUOR 340) objective lenses equipped with an immersion corrected dipping cap mounted on a LaVision UltraII microscope coupled to a white light laser module (NKT SuperK Extreme EXW-12) for imaging. The images were taken in 16 bit depth and at a nominal resolution of 3.25 µm and 1.625 µm / voxel on the XY axes (for the 2x and 4x respectively). The brain vasculature was visualized by DyLight 594 (using a 580/25 nm excitation and a 625/30 nm emission filter). To reduce defocus, which derives from the Gaussian shape of the beam, we used a 5-step sequential shifting of the focal position of the light-sheet per plane and side. In z-dimension we took the sectional images in 32.5 µm and 3 µm steps (for the 2x and 4x respectively) using left and right sided illumination. Perfused brain vasculature was detected and quantified using our previously established Vessel Segmentation and Analysis Pipeline (VesSAP) from the 4x scans [15]. Quantitative metrics included regional vessel length normalized to brain region volume, bifurcation density (number of branch points normalized to region volume), and the radius of the vessels. All measurements were corrected by a constant factor to account for tissue shrinkage introduced by fixation and clearing. 3D renderings were made with Imaris 9.9 (Oxford Instruments).

**Statistics.** Statistical analysis was performed in GraphPrism. Two-tailed Student's t-test was used for the comparison of two groups. If variances were unequal, Welch correction was used. In case a Gaussian distribution could not be assumed (e.g. due to a floor effect while reducing already low gene expression to almost zero), the Mann-Whitney-test was used. For the combined data set of both sexes, samples of each sex were normalized to the respective controls to set all controls to an average of 1. One-way ANOVA and Tukey *post hoc* test were used when one factor was varied. Two-way ANOVA and Bonferroni *post hoc* test were used when two factors were varied. The cumulative head diameter of dendritic spines was analyzed using the Gehan-Breslow-Wilcoxon test [16]. Outliers were removed when the data points were more than two standard deviations away from the average. An exception was the postmortem analysis, where samples were excluded before analysis based on extreme age or postmortem interval values to avoid age/postmortem biases between data sets. Most experiments were performed only once to minimize animal numbers and reduce variability. An exception was behavioral testing, which generally requires larger group sizes. To further increase robustness, a variety of tests and measures was applied (e.g. various behavioral tests, morphology, molecular analysis). Moreover, certain conditions such as CVS were performed in more than one experiment, leading to partial replication.

Transcriptomics, proteomics, dendritic spine analysis and brain vasculature measurements were each performed in different cohorts of animals. Each of these experiments was analysed by different experimenters. Behavioral, dendritic and vasculature analyses were performed blinded to the condition. Transcriptomics and proteomics were measured by two different core facilities. Hence, the chance of bias across this paper was minimized as much as possible.

Heatmaps depict significant gene products of the upper lane, sorted by the log2 fold change. In the lower lane, all corresponding gene products of compared group are shown, regardless of their significance. Venn diagrams show significant gene products of both groups. Sizes of the circles are not in scale. Metascapes were generated as described [17].

To assess the degree of overlap in protein changes across multiple datasets, the improved rank–rank hypergeometric overlap (RRHO) test was used [18,19]. Gene products were ranked according to their adjusted P-values with ranks assigned directionality according to fold-change values. Pairwise comparisons between ranked lists were conducted using hypergeometric testing, generating a two-dimensional matrix of p-values. Multiple testing was controlled using the Benjamini–Yekutieli false discovery rate correction, and adjusted p-values were visualized as heatmaps.
